# Phenotypic analysis of catastrophic childhood epilepsy genes: The Epilepsy Zebrafish Project

**DOI:** 10.1101/2021.02.11.430844

**Authors:** Aliesha Griffin, Colleen Carpenter, Jing Liu, Rosalia Paterno, Brian Grone, Kyla Hamling, Maia Moog, Matthew T. Dinday, Francisco Figueroa, Mana Anvar, Chinwendu Ononuju, Tony Qu, Scott C. Baraban

**Author notes:** These authors contributed equally to this work (Aliesha Griffin, Colleen Carpenter). Correspondence should be addressed to, Correspondence and requests for materials should be addressed to Scott C. Baraban.

## Abstract

Genetic engineering techniques have contributed to the now widespread use of zebrafish to investigate gene function, but zebrafish-based human disease studies, and particularly for neurological disorders, are limited. Here we used CRISPR-Cas9 to generate 40 single-gene mutant zebrafish lines representing catastrophic childhood epilepsies. We evaluated larval phenotypes using electrophysiological, behavioral, neuro-anatomical, survival and pharmacological assays. Phenotypes with unprovoked electrographic seizure activity (i.e., epilepsy) were identified in zebrafish lines for 8 genes; *ARX, EEF1A, GABRB3, GRIN1, PNPO, SCN1A, STRADA* and *STXBP1*. A unifying epilepsy classification scheme was developed based on local field potential recordings and blinded scoring from ~3300 larvae. We also created an open-source database containing sequencing information, survival curves, behavioral profiles and representative electrophysiology data. We offer all zebrafish lines as a resource to the neuroscience community and envision them as a starting point for further functional analysis and/or identification of new therapies.

## Introduction

Catastrophic childhood epilepsies are characterized by intractable persistent seizures and are frequently associated with developmental delay, cognitive dysfunction and autism^1–3^. Many are rare genetic disorders lacking effective therapeutic options^4–6^. With technological advances and large-scale patient cohorts, genome-wide analyses have now identified *de novo* mutation in a single gene for most of these epilepsies^7–11^. These studies highlight the complexity of epilepsy, as mutations in genes coding for ion channels, ligand-gated receptors, solute transporters, metabolic enzymes, synaptic trafficking proteins, kinases, transcription factors, and adhesion molecules were identified. Unfortunately, our overall understanding of genetic epilepsies is severely limited as few experimental animal models exist, and human induced pluripotent stem cell derived two- or three-dimensional neuronal models fail to fully recapitulate the complex brain network seen in patients. Zebrafish, a small vertebrate with considerable genetic similarity to humans^12^, offer an attractive alternative model to study these genetic mutations *in vivo*. Analysis of zebrafish mutants for human genes has provided valuable insight into complex circuits controlling behavior^13–17^, evolutionarily conserved developmental programs^18–20^ and drug candidates for a variety of diseases, including epilepsy^21–29^.

Epilepsy classification, incorporating an understanding of different seizure types and comorbidities, is an essential clinical resource in evaluating patients and selection of anti-seizure treatments^30–33^. Clinical classification resources have evolved continuously since the 1960s. However, adaptation of this classification strategy to animal models^34^, specifically zebrafish models developed for catastrophic epilepsies of childhood, is lacking. Because clinical seizure classifications promoted by the International League Against Epilepsy (ILAE)^33^ are defined by the presence of unprovoked “self-sustained paroxysmal disorders of brain function”, we focused our phenotyping effort on developing a standardized seizure classification scheme using electrophysiology data. Such a resource, broadly adapted, could be particularly useful for preclinical studies designed to characterize epilepsy phenotypes in any larval zebrafish model.

To better understand mechanisms underlying human genetic epilepsies, it is important to first identify clinically relevant phenotypes in an experimental model system^35^. Although efficient gene inactivation in mice has contributed many pediatric epilepsy models^36–38^, to generate dozens of mutant mouse lines followed by a systematic phenotypic analysis would require several decades of research. Using an efficient CRISPR-based gene editing strategy^39,40^ we successfully generated 37 stable zebrafish lines representing human monogenic pediatric epilepsies. Large-scale phenotypic analysis of survival, behavior and electrographic brain activity was performed. We established read-outs to identify seizures at electrographic and behavioral levels, and an open-source online website to efficiently share data with the neuroscience community. As many of these zebrafish represent rare genetic diseases for which our understanding of pathophysiology remains largely unknown, they provide a rich resource to further investigate key etiological questions or utilization in high-throughput precision medicine-based therapy development.

## Results

### Generation of loss-of-function models for human epilepsy genes

We evaluated genes identified in a genome-wide association study from 264 patients with epileptic encephalopathies by the world-wide Epilepsy Genetics Initiative, Epi4K Consortium^8,41^. First, analysis of human genetic data was performed to identify genes where a loss-of-function (LOF) mutation was likely a causal mechanism of the epileptic phenotype. This limited our initial Epilepsy Zebrafish Project (EZP) choices to 63 gene candidates (Supplementary Table 1). Second, Epilepsy Genetics Initiative identified human genes were selected representing 57 orthologous zebrafish genes (Figure 1a). From this group, we identified 48 zebrafish genes that were high confidence orthologs (Figure 1b, homology scores; Supplementary Table 2) and examined expression data patterns with a primary focus on brain expression (Figure 1c). Third, RT-PCR confirmed gene expression for 46 zebrafish orthologs from the 4-cell to 7 dpf stage (Figure 1e) e.g., an early neurodevelopmental window wherein high-throughput studies would be feasible. To generate stable mutant lines, we used Cas9 with single *in vitro* transcribed guide RNA (with no predicted off-target sites) targeted towards the start of the protein coding sequence. A total of 46 zebrafish orthologous genes were targeted (Supplementary Table 3). This group includes a previously published *stxbp1b* mutant^42^ and a novel *scn1lab* CRISPR mutant. Adult founders harboring predicted protein coding deletions (Figure 1e; https://zebrafishproject.ucsf.edu) were confirmed and outcrossed for at least two generations. All EZP zebrafish were maintained as outcrossed lines with phenotypic assessment(s) performed on larvae generated from a heterozygous in-cross. For seven genes we could not obtain a viable line (*grin2aa*, *syngap1a*, *tbc1d24, prickl1a*, *plcb1*, *gosr2* and *stx1b*). In total, 37 novel EZP zebrafish lines were subjected to phenotypic screening described below.

**Figure 1.**
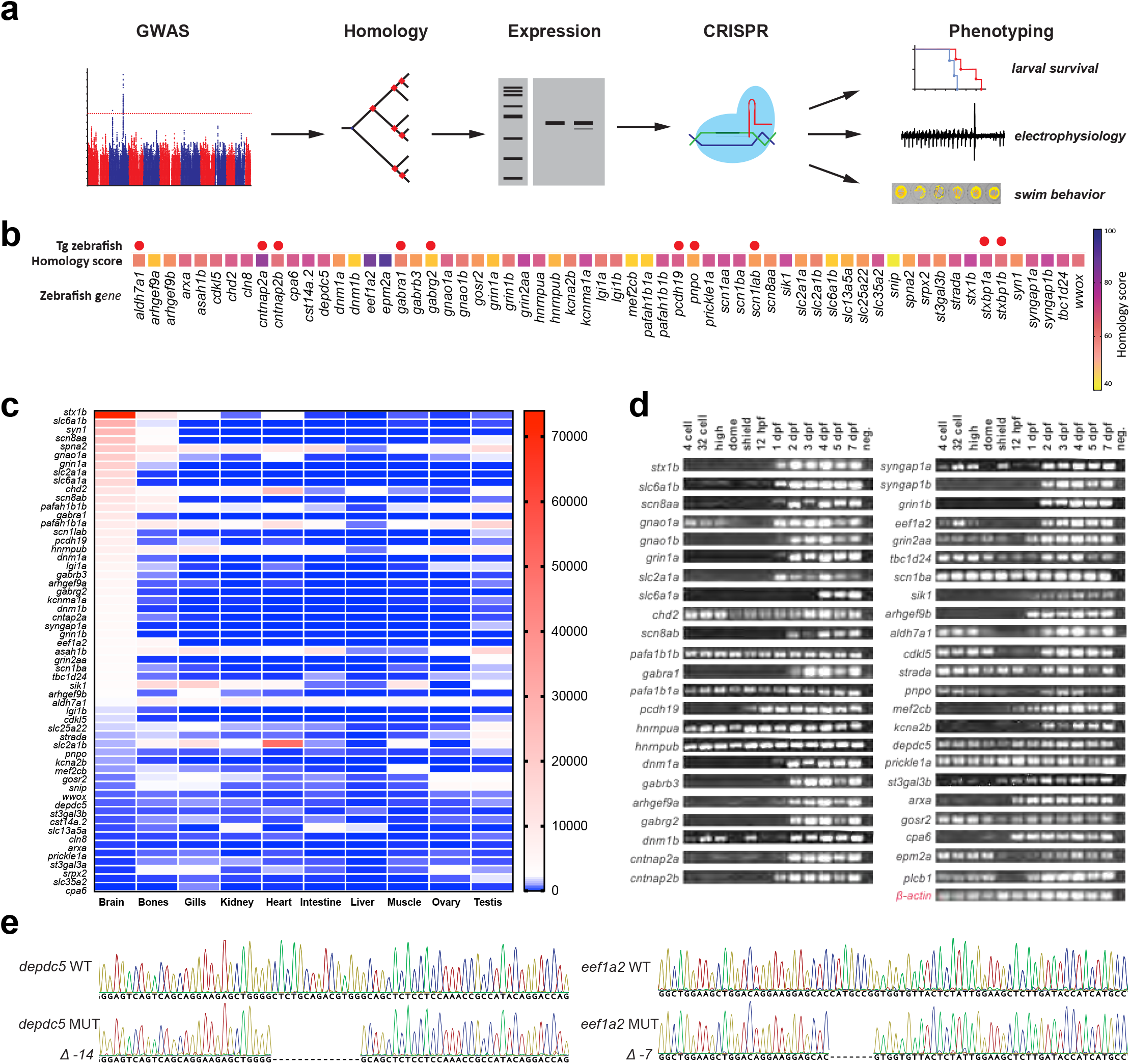
The Epilepsy Zebrafish Project (EZP). (**a**) Overview of the zebrafish epilepsy disease model discovery workflow from human genome wide association studies (GWAS) to generation of zebrafish models and phenotypic characterization. (**b**) Tissue expression profiles of EZP zebrafish target genes. Heatmap represents the maximum number of sequence reads for each gene per tissue. (**c**) Developmental gene expression profiles for EZP lines. (**d**) Representative frame-shift mutant lines confirmed for *depdc5* and *eef1a2*.

### Classification of seizure activity in larval zebrafish

We previously described minimally invasive local field potential recording (LFP) techniques to monitor brain activity in larval zebrafish^43^ (Supplementary Figure 1). To identify epilepsy phenotypes in CRISPR-generated zebrafish lines, we obtained LFP recordings from 3255 larvae at 5 and 6 days post fertilization (dpf). We blindly recorded a minimum of 75 larvae per line, from at least three independent clutches. Larvae were randomly selected and genotyped *post hoc* to evaluate homozygote, heterozygote and wild-type (WT) phenotype-genotype correlations. Although long-duration, multi-spike large amplitude discharges are commonly described as seizure events in larval zebrafish models^23,44–47^, a unified seizure classification system does not exist. As seizure classification is an essential clinical tool in identification of an epilepsy phenotype^48^, we sought to establish the first classification scheme that could be universally applied to all zebrafish epilepsy models. An LFP electrophysiology-based scoring system covering all types of observed activity was established: (i) **Type 0**: the range of low voltage activities and patterns of small membrane fluctuations; (ii) **Type I**: low amplitude *interictal-like* sharp waveforms, with voltage deflections at least three times above baseline (duration range: 10 - 99 msec); and (iii) **Type II**: large amplitude *ictal-like* multi-spike waveforms, with voltage deflections at least five times above baseline (duration range: 45 - 5090 msec), often followed by a transient period of electrical suppression with no detectable events (Figure 2a). Based on this numeric classification, each 15 min recording epoch was assigned an LFP score by two independent investigators; cumulative averages can be seen in the heatmap for all 37 EZP-generated zebrafish lines (Figure 2b). We classified mutants with an average LFP score of 1.0 or above as an epilepsy phenotype. These included two genes previously determined to exhibit epilepsy phenotypes in zebrafish (e.g., *scn1lab^23^* and *stxbp1b* homozygotes^42^) and six novel zebrafish epilepsy lines (e.g., *arxa*, *eef1a2*, *gabrb3*, *pnpo, strada* homozygotes and *grin1b* heterozygotes). The percentage of EZP mutant larvae scored at Type II ranged from 29 to 83% for epilepsy lines and a significant correlation between LFP classification scores versus percentage of Type II mutants was noted (Figure 2c; R^2^ = 0.8790). Distribution of LFP classification scores for all WT larvae skewed toward Type 0 (mean WT score = 0.66; n = 781) and was significantly different than scoring distributions for mutant lines designated as epileptic (mean EZP-epilepsy score = 1.23; n = 190; Unpaired t-test *p* < 0.0001, t = 10.26, df = 969)(Figure 2d). The majority of LFP recordings from all lines were classified as Type 0 or 1 (79%; n = 3255; Figure 2e).

**Figure 2.**
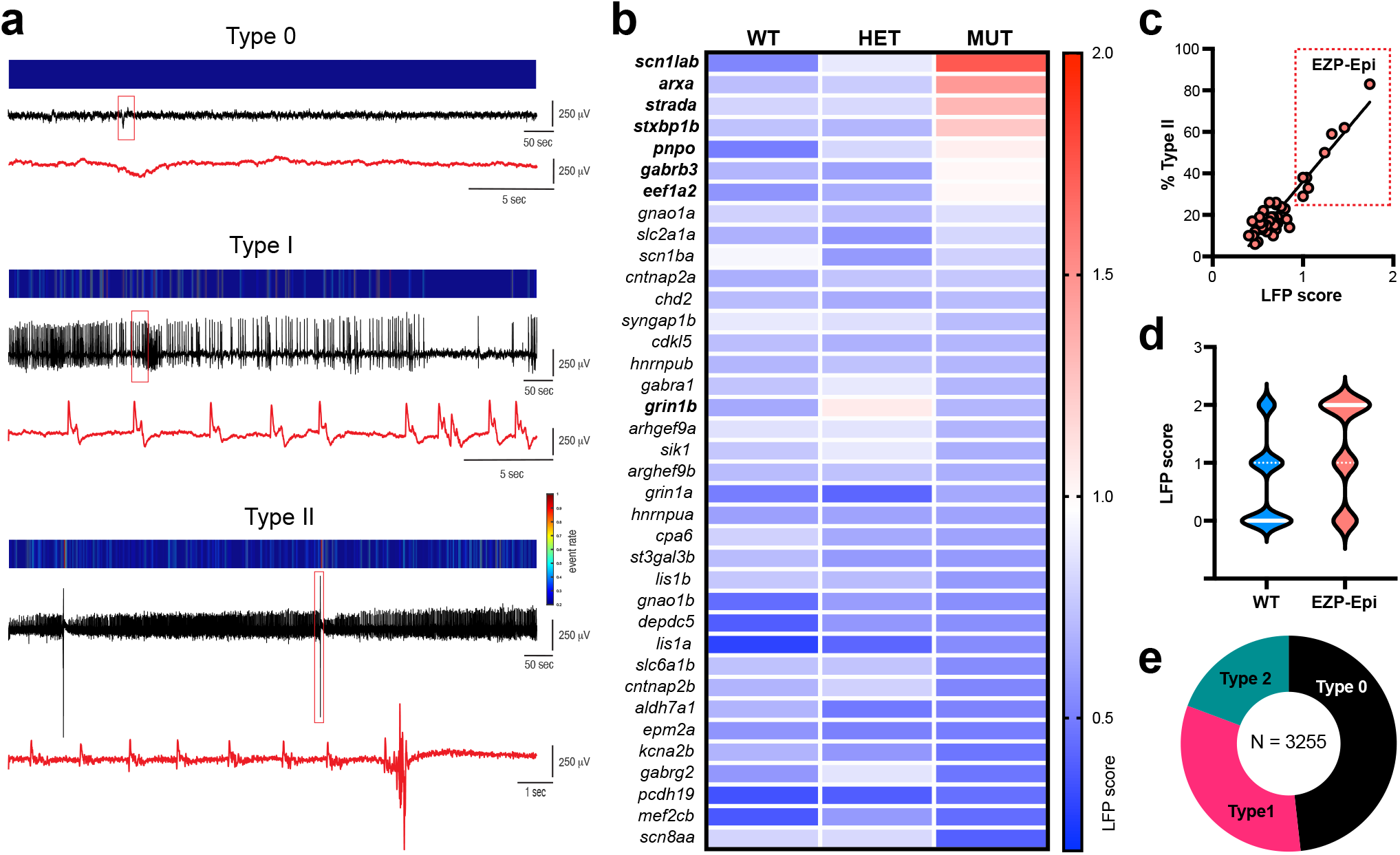
Seizure classification using electrophysiological recording identifies epileptic zebrafish lines. (**a**) LFP recordings representing Type 0 (low voltage, small or no membrane fluctuations), Type I (low amplitude, sharp *interictal-like* waveforms) and Type II (low frequency, sharp *ictal-like* waveforms with large-amplitude multi-spike events and post-ictal slowing) scoring activity. For each example a color-coded event rate histogram (top), full 15 min LFP recording (middle), and high-resolution LFP close-up (red box, red trace at bottom) are shown. (**b**) Heatmap showing mean larval zebrafish LFP recording scores for all 37 EZP zebrafish lines ranked from highest homozygote score to lowest; N = 77 to 127 larvae per gene (see https://zebrafishproject.ucsf.edu for N values on each individual line). A threshold of a mean LFP score > 1.0 was classified as a an EZP line exhibiting epilepsy (indicated in bold font: *scn1lab*, *arxa*, *strada*, *stxbp1b*, *pnpo*, *gabrb3*, *eef1a2* and *grin1b*). (**c**) Regression plot for all 37 mutants showing mean LFP score versus % of Type II larvae for each homozygote. 7 homozygote and 1 heterozygote lines highlighted in “EZP-epi” box as clearly differentiated from cluster of 31 non-epileptic EZP lines with LFP scores < 1.0. Simple linear regression R^2^ = 0.8790; ***Significant deviation from zero, p < 0.0001; DFn, DFd = 1, 36. (**d**) Violin plots of all LFP scores recorded for EZP-epilepsy lines (N = 190) compared to all WT control siblings (N = 783). Note: Type 2 epileptiform events were only observed in 14.7% of all WT larvae. (**e**) Distribution of Type 0, I and II scores for all WT, heterozygote and homozygote larvae screened by LFP recordings (N = 3255).

We next examined the frequencies, durations and spectral features of spontaneous epileptiform events recorded in all 8 EZP-epilepsy lines. To provide an unbiased quantitative analysis, Type I interictal- and Type II ictal-like electrical events were detected using custom software (see Methods; Figure 3) on homozygote and WT sibling larvae recordings. Representative LFP recordings (Figure 4b, top) with accompanying time-frequency spectrograms (Figure 4b, bottom) are shown for each EZP epilepsy line; individual LFP scoring distribution plots for mutants and WT siblings are shown at left. No difference in interictal-like (Type I) event frequency or duration was noted (Figure 4c). Ictal (Type II) events were more frequent and longer in duration for *scn1lab* mutant compared to WT; ictal event duration was shorter for *stxbp1b* mutants compared to WT (Figure 4d). Ictal event histograms showed similar overall distributions at a cumulative and individual level (Figure 5; Supplementary Figure 2). However, large-amplitude multi-spike ictal events when present in WT siblings were usually brief in duration, rarely exceeding 2.0 sec (Figures 5a, 5c) and less frequently encountered (Figures 5b, 5c) than those identified in EZP-epilepsy lines (also see cumulative distribution insets in Supplementary Figure 2a). Representative raw LFP traces and classification distribution plots for all 37 zebrafish lines can be explored on our open-source website, https://zebrafishproject.ucsf.edu, where users can also find information on homology, sequencing, survival and genotyping protocols.

**Figure 3.**
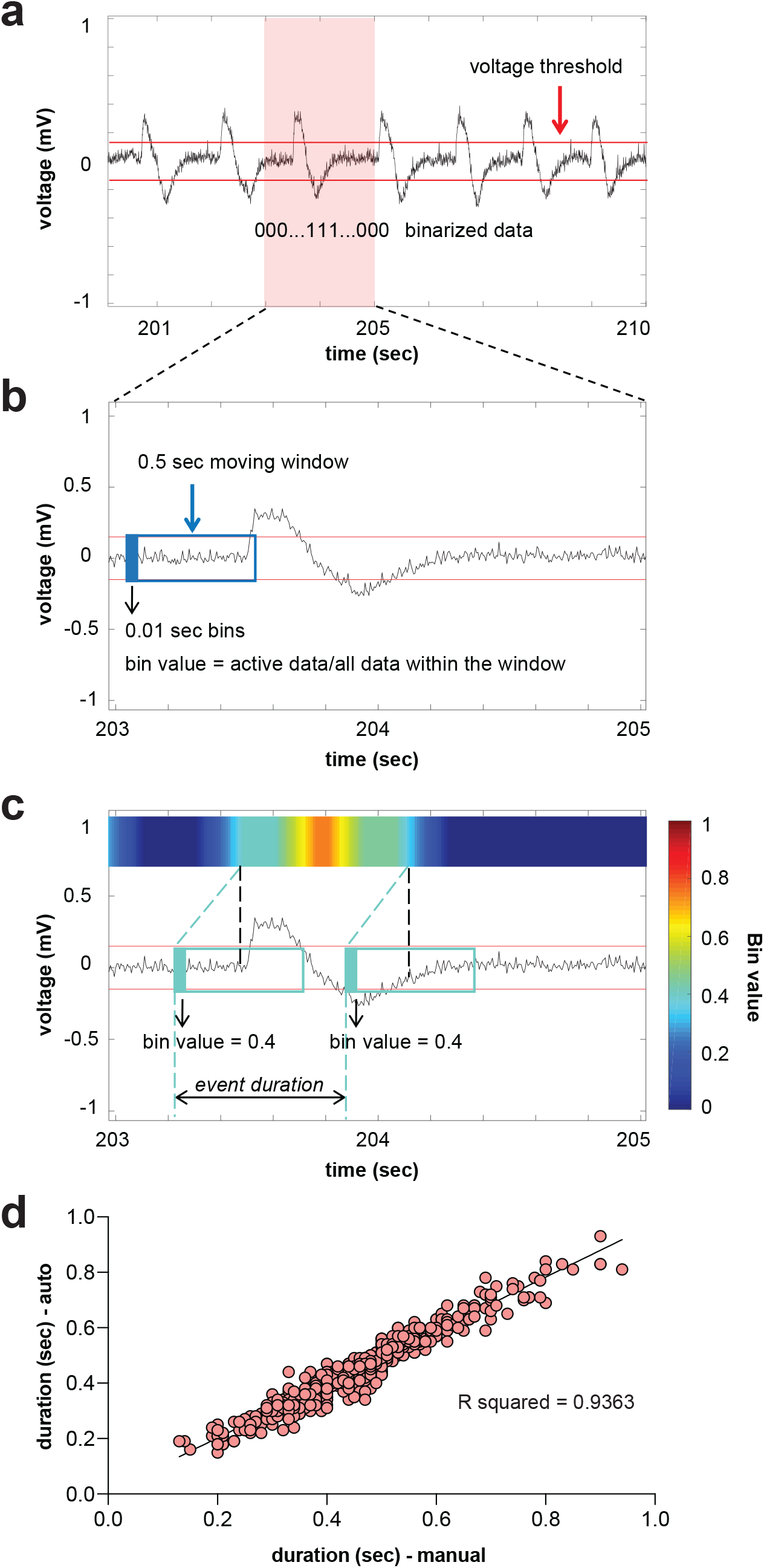
Automated interictal-like event quantification. (**a**) A representative LFP recording with interictal-like events. A voltage threshold (0.15 – 0.25 mV, depending on the noise level) was set for event detection. Data was binarized by threshold: super-threshold data points were scored as 1, and under-threshold data points were scored as 0. (**b**) A data binning method was used for automated quantification of interictal-like events: 0.01 sec binning in 0.5 sec time window. In each window, value of the first bin was calculated, which is the ratio of active data points to the number of total data points within the window. (**c**) Color raster plots were created according to the raster score. A raster score threshold (0.2 – 0.4) was set to define the start and end of an event. (**d**) Comparison between interictal-like event durations measured automatically and manually. A 10 sec representative epoch from each recording will be used as a testing sample to optimize the algorithm. Voltage and raster score thresholds were chosen when the difference between automated and manual results is less than 3% of manual measurements.

**Figure 4.**
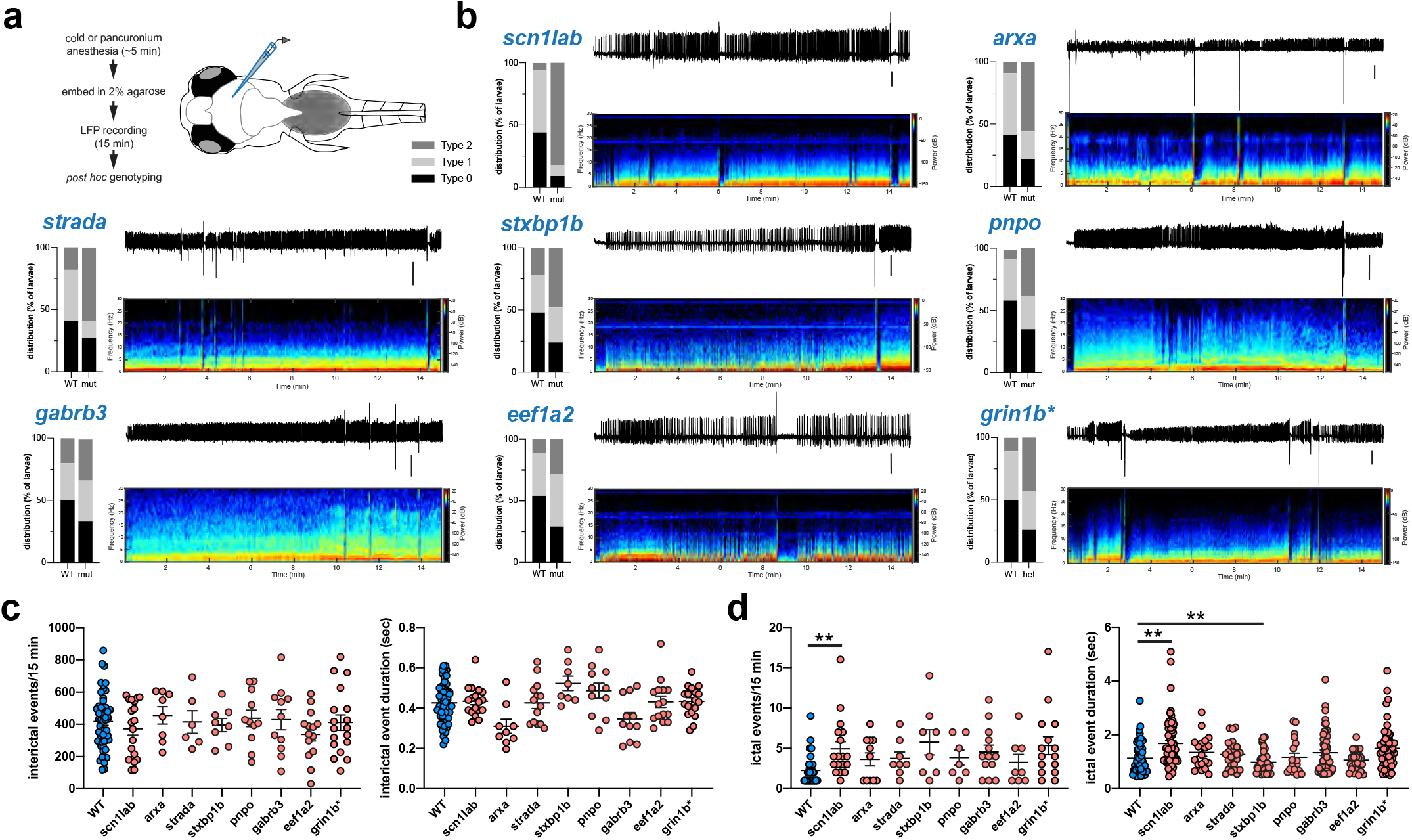
Electrographic seizure activity in epileptic zebrafish mutant lines. (**a**) Schematic of recording configuration and protocol for electrophysiology-based screening of larval zebrafish. (**b**) Representative raw LFP recording traces (top, right) along with a corresponding wavelet time-frequency spectrogram (bottom, right) and LFP scoring distribution plot for WT and mutant larvae (left) are shown for each EZP-epilepsy line. Type 0, I and II scoring as in Figure 2. A representative WT LFP recording with the corresponding wavelet time-frequency spectrogram is shown in Supplementary Figure 5. Scale bar = 500 μV. Representative LFP recordings and distribution plots for all 37 lines can be found online (https://zebrafishproject.ucsf.edu). (**c**) Cumulative plots of interictal event frequency (left) and duration (right) for all EZP-epilepsy lines compared to WT sibling controls. Each point represents mean of all interictal events in a single 15 min larval LFP recording detected using custom software in MATLAB (N = 9775, WT; N = 6750, *scn1lab*; N = 2550, *arxa*; N = 5790, *strada*; N = 6750, *stxbp1b*; N = 3538, *pnpo*; N = 3455, *gabrb3*; N = 4335, *eef1a2*; N = 6610, *grin1b*\*). (**d**) Cumulative plots of ictal event frequency (left) and duration (right). Each point represents all ictal events in a single 15 min larval LFP recording detected using custom software in MATLAB (N = 56, WT; N = 62, *scn1lab*; N = 26, *arxa*; N = 26, *strada*; N = 48, *stxbp1b*; N = 22, *pnpo*; N = 59, *gabrb3*; N = 27, *eef1a2*; N = 55, *grin1b*\*). *for *grin1b* designates heterozygote. \*\**p* < 0.01, ANOVA with Dunnett’s multiple comparisons test.

**Figure 5.**
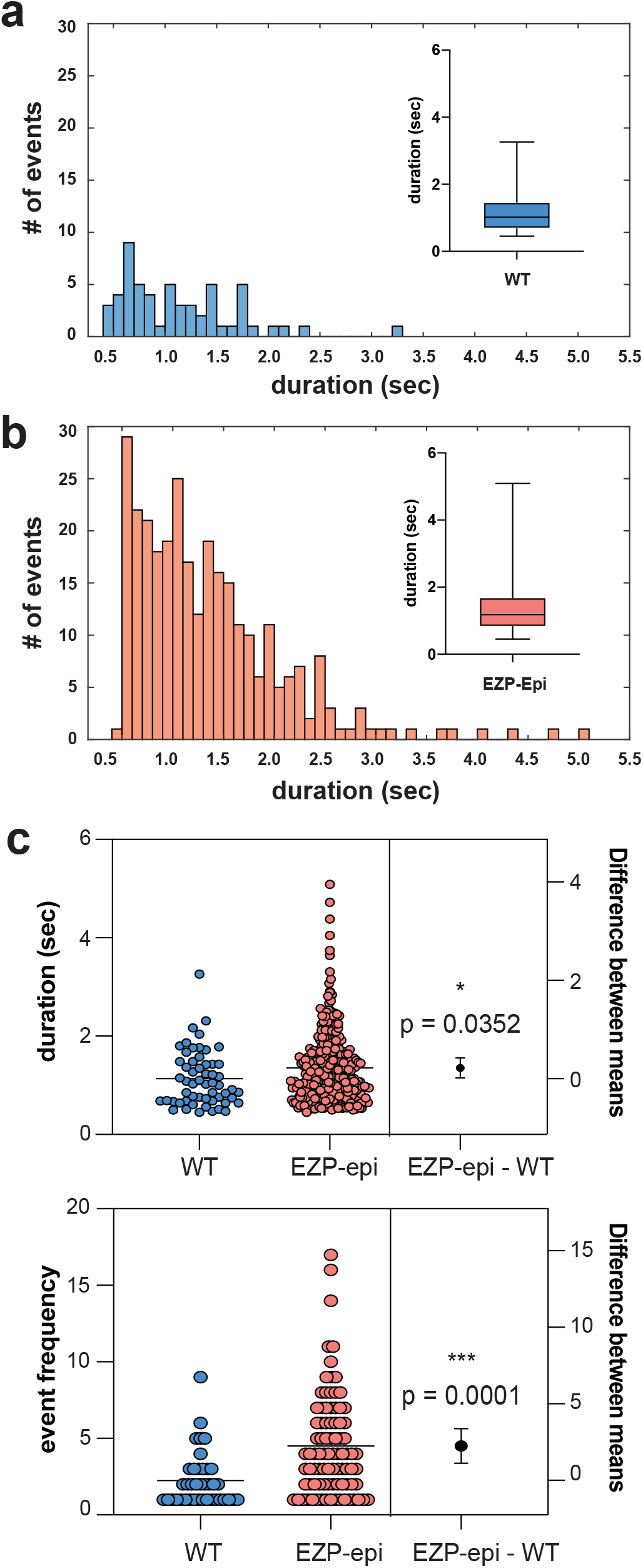
Distribution of ictal events. Histograms depict number and duration of ictal events measured using a custom MATLAB-based program for (**a**) all sibling wild-type (WT) larvae from EZP epilepsy lines and (**b**) same for epileptic zebrafish lines (EZP)-Epi. Box-and-whisker plots showing the distribution of ictal event durations; mean and minimum/maximum values are shown (*insets*). (**c**) Estimation plot showing that ictal event duration for WT (1.134 ± 0.075 sec; N = 56) is shorter than for Epi-EZP (1.353 ± 0.043 sec; N = 299); Non-parametic t-test \**p* = 0.0352, t = 2.115, df = 353). Each dot on the top plot represents the duration (measured in msec) for one individual ictal event.; each dot in the bottom plot represents ictal event frequency for one LFP recording. LFP recording epochs were 15 min.

### EZP lines for understanding disease pathophysiology

Epileptic zebrafish can be used to study underlying neurobiological mechanisms, behavioral comorbidities and drug discovery. Many pediatric epilepsies are associated with increased mortality rates and thus, survival studies were performed on all EZP lines to evaluate larval health, confirm Mendelian genotyping ratios, and identify early death phenotypes (Figure 6a, Supplementary Figure 3). Early fatality was noted in *aldh7a1*, *depdc5*, *scn8aa* and *strada* homozygous mutants that only survive between 8-10 dpf (Figure 6b).

**Figure 6.**
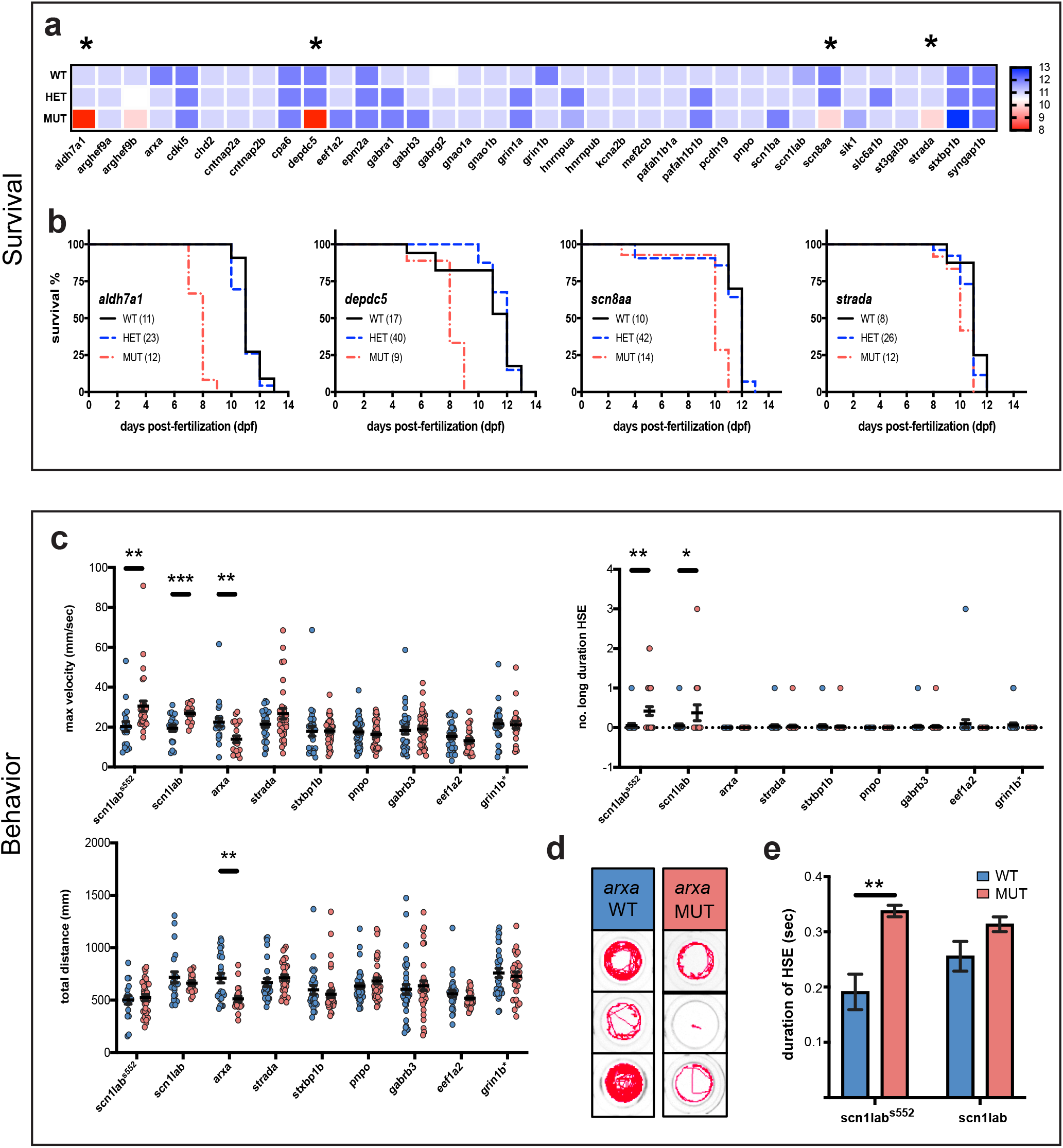
Survival and behavioral phenotypes. (**a**) Heatmap displaying median wild-type (WT), heterozygote (HET) and homozygote mutant (MUT) larval survival for EZP lines. Range extends from 8 dpf (red) to 13 dpf (blue). Asterisks indicate MUTs with significant survival deficits compared WT control siblings; p < 0.05, log rank test. (**b**) Lines with significant survival deficits. (**c**) Quantification of the basal locomotor activity of epileptic lines after 1 hr habituation in DanioVision chamber. Maximum velocity and total distance traveled were extracted directly from EthoVision XT 11.5 software while the number of events ≥ 28 mm/s, termed high speed events (HSE), and long duration HSE (≥ 1 s) were scored using a MATLAB algorithm (*scn1lab^552^* WT N = 19, MUT N = 31; *scn1lab* WT N = 21, MUT N = 16; *arxa* WT N = 25, MUT N = 22; *strada* WT N = 27, MUT N = 31; *stxbp1b* WT N = 26, MUT N = 43; *pnpo* WT N = 42, MUT N=40; *gabrb3* WT N = 35, MUT N = 36; *eef1a2* WT N = 30, MUT N = 27 and *grin1b* WT N=29 and HET=57). (**d)** Representative traces of *arxa* WT and MUT movement. (**d**) Comparison of duration of HSE in *scn1lab* ENU and CRISPR larvae. Displayed as mean ± SEM, One-Way ANOVA was used to determine the significance of both HET and MUT behavior for all lines (See Supplementary Figure 4 for expanded data set). *Post hoc* Dunnett multiple comparison test, \**p* ≤ 0.05, \*\**p*≤ 0.005, \*\**p* < 0.0001.

We further performed a series of pilot experiments in all 8 EZP-epileptic lines to investigate other known pathophysiology. Epilepsy often manifests as convulsive behaviors in many of these genetic epilepsies. Prior work from our laboratory using chemically induced (Pentylenetetrazole ; PTZ) or an ENU-mutagenesis mutant for Dravet syndrome (*scn1lab^s552/s552^*) describe a characteristic series of larval seizure-like behaviors, culminating in bursts of high-speed swim activity and whole-body convulsions^23,44^. Using these well-established models, we first developed a custom MATLAB algorithm to detect high-speed (≥ 28 mm/s), long-duration (≥ 1 s) behavioral events corresponding to these convulsive behaviors in freely behaving larvae (Figure 7). The MATLAB-detected behavioral event duration was similar to that measured for Type II ictal-like events in LFP recordings (see Figure 5). As expected, the EZP generated *scn1lab* mutant larvae displayed significantly higher velocity movements and higher frequencies of convulsive-like events compared to WT sibling controls; similar results were obtained with *scn1lab^s552/s552^* larvae. There was no difference in the total distance traveled between WT and homozygous mutants in these lines (Figure 6c). Maximum velocity and total distance measurements show that *arxa* larvae are hypoactive and they had no detectable high-speed, long-duration events during these 15 min recording epochs (Figure 6c; Figure 6d, representative traces). We observed that the duration of high speed events in *scn1lab^s552/s552^* larvae were significantly longer than in WT sibling controls (Figure 6e). No significant behavioral phenotypes were seen in the other epileptic lines (Supplementary Figure 4).

**Figure 7.**
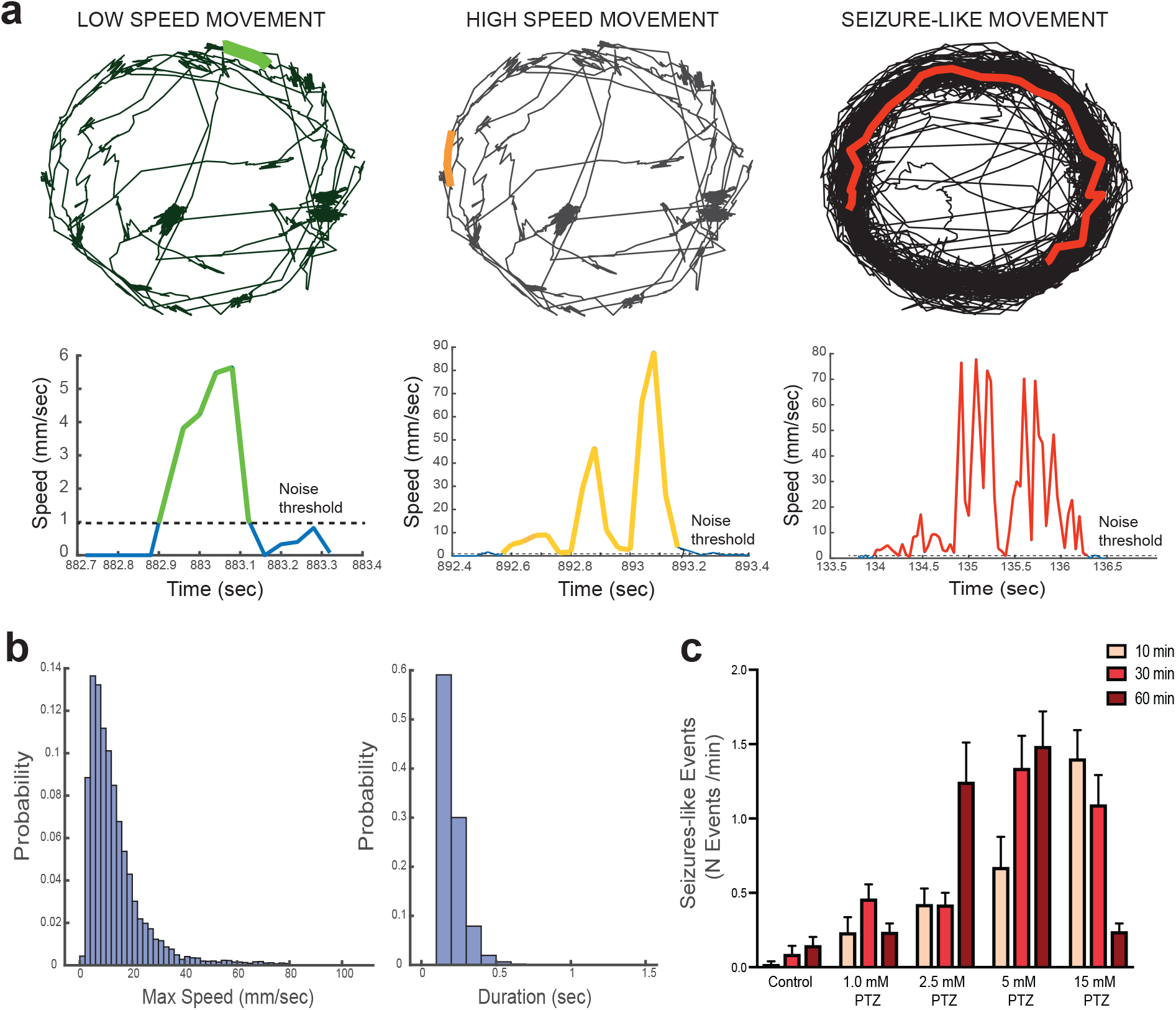
Automated detection of behavioral seizure-like events. (**a**) Example of low-speed movement in a WT larva (left - green), high-speed movement in the same WT larva (middle - orange), and seizure-like movement in a PTZ-treated larva (right - red). Top traces represent the larvae track during 15 min recording in a 96-well plate. The bottom panels show speed values across time for the events highlighted. Note the short and long duration in the high-speed events in WT and PTZ-treated larvae, respectively. (**b**). Distribution of maximum speed (left) and duration (right) across all movements in WT larvae (n: 109) during the 15-minute recording session. The average maximum speed was 10.5 mm/sec and the duration of the events was less than 1 second. (**c**). Frequency of seizure-like movements (defined as events with maximum speed greater than 28 mm/sec and duration greater than 1 second) in control and PTZ-treated larvae at different concentrations after 10, 30 and 60 minutes (two-way ANOVA p<0.05). Note the increased number of events with increasing PTZ dose and the lower number when using 15 mM after 60 minutes due to increased larvae mortality.

*ARX*-related epilepsies are categorized as “interneuronopathies”^49^ and *Arx* mutant mice exhibit a reduced number of interneurons in both neocortex and hippocampus^50,51^. Using volumetric light-sheet microscopy imaging in larval *arxa* mutants co-expressing a green fluorescent protein (GFP) in *Dlx*-labeled interneurons^52^, we confirmed a significant reduction in interneuron density for homozygous *arxa* mutant larvae compared to WT sibling controls (Figure 8a). *EEF1A2* mutations are associated with neurodevelopmental deficits in some patients.^53^ Using conventional morphological analyses measuring overall head length, midbrain/forebrain width and body length on *in vivo* images from *eefla2* mutant larvae and WT siblings at 5 dpf, we noted no differences (Figure 8b).

**Figure 8.**
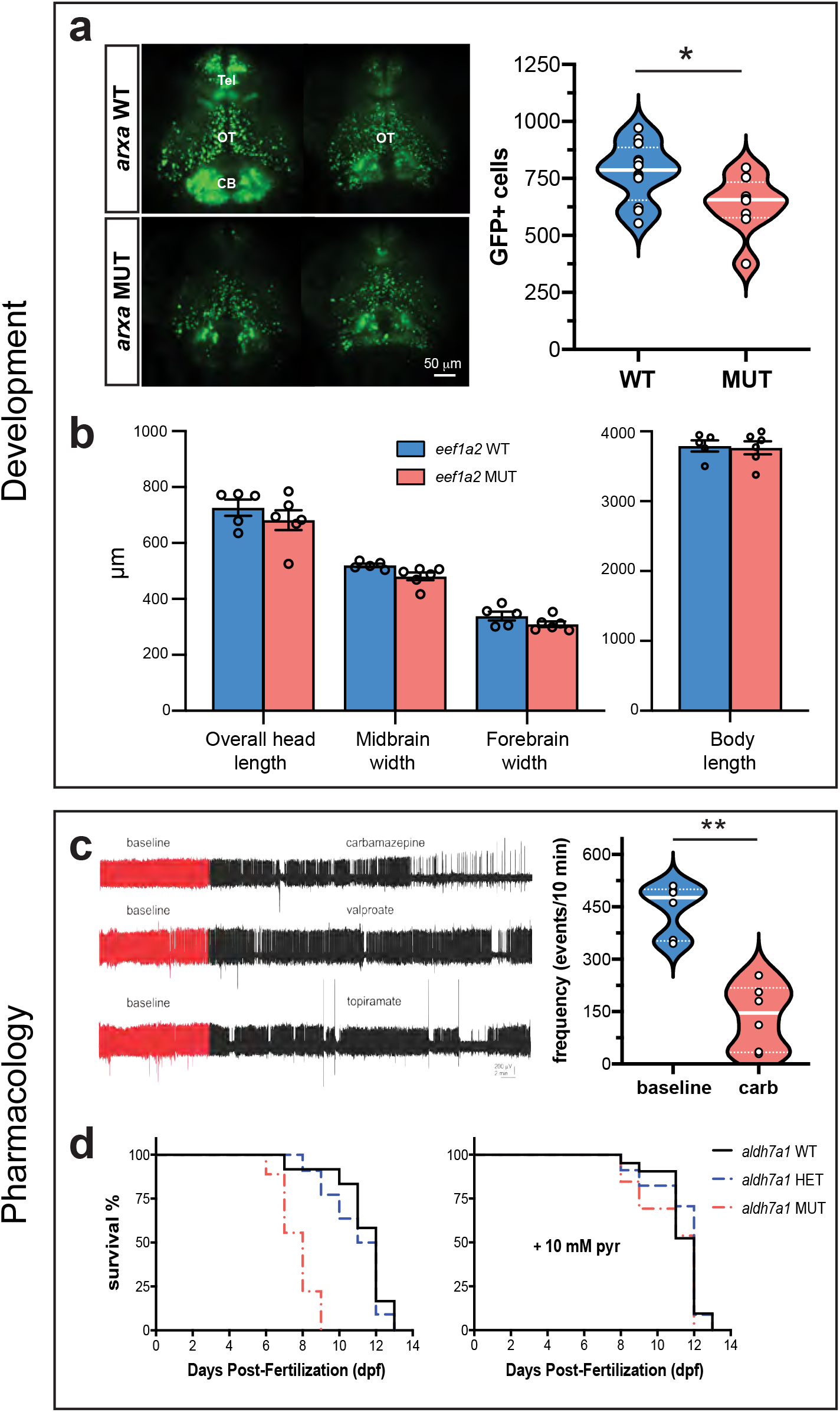
Developmental and pharmacological characterization. (**a**) Representative images of *dlx*-GFP expressing interneurons in *arxa* MUT larvae (N = 8) and WT siblings (N = 12) obtained from volumetric light sheet imaging microscopy. Unpaired two-tailed t-test \**p* = 0.0268; t = 2.411, df = 18) (**b**) High resolution images of larvae were taken using a SteREO Discovery.V8 microscope (Zeiss) and overall head length, midbrain width, forebrain width and body length were quantified in *eef1a2* MUT (N=6) and WT (N=5) larvae. (**c**) Representative 1 hr LFP traces from *gabrb3* MUT larvae exposed to AEDs. The first ~10 min of the recording (in red) represents baseline. Drugs were bath applied at a concentration of 0.5 mM; N = 3-6 fish per drug. Results from carbamazepine treatment shown as violin plot. Unpaired two-tailed t-test \*\**p* < 0.0001; t = 6.344, df = 10). (**d**) Kaplan-Meier survival curves for *aldh7a1* WT, *aldh7a1* HET and *aldh7a1* MUT larvae treatment with 10 mM pyrodixine (pyr) or vehicle for 30 mins daily starting at 4 dpf. Median survival for vehicle treated *aldh7a1* WT *=* 12 dpf (N = 12), *aldh7a1* HET = 11.5 dpf (N = 22) and *aldh7a1* MUT = 8 dpf (N = 9). Median survival for 10 mM pyridoxine (pyr) treated larvae for *aldh7a1* WT *=* 12 dpf (N = 21), *aldh7a1* HET = 12 dpf (N = 34) and *aldh7a1* MUT = 12 dpf (N = 13).

Patients with *GABRB3* mutations, like many of the genes studied here, are often classified as pharmaco-resistant^54^. Using a 1 hr LFP recording protocol, we evaluated electrographic seizure activity in *gabrb3* mutants treated with standard antiepileptic drugs (AEDs): carbamazepine, valproate and topiramate. In these zebrafish mutants, carbamazepine suppressed high-frequency interictal-like and long duration multi-spike ictal-like epileptiform discharges (Figure 8c). Patients with *ALDH7A1* mutations are associated with pyridoxine-dependent encephalopathy. Using a CRISPR-generated *aldh7a1^ot100^* mutant, Pena et al. reported hyperactive behavior and spontaneous electrographic seizures in fed larvae starting at 10 dpf; 10 mM pyridoxine treatment rescued these phenotypes^46^. The unperturbed EZP generated *aldh7a1* mutant larvae die prematurely between 7 and 9 dpf. Daily 10 mM pyridoxine effectively extended the median survival of *aldh7a1* mutant larvae to that observed in heterozygote and WT sibling controls (Figure 8d).

## Discussion

Progress in exploring pathogenesis, and developing new therapies, for monogenic epilepsies is complicated by limited availability of preclinical animal models for many of these genes. The emergence of zebrafish as a vertebrate model system amenable to genetic manipulation holds much promise toward accelerating progress in understanding these rare epilepsies. Here we utilized CRISPR/Cas9 and a battery of larval zebrafish assays to systematically evaluate 40 different single gene mutations identified in this population. We determined that homozygous deletion of *arxa*, *eef1a2*, *gabrb3*, *pnpo*, *scn1lab*, *strada* and *stxbp1b* or heterozygous loss of *grin1b* result in recurrent unprovoked electrographic seizures (i.e., epilepsy). In addition, we developed an electrophysiology-based classification system that can be used to identify seizures in any larval zebrafish model. Finally, we show that clinically relevant phenotypes such as interneuron loss (*arxa*) or pharmaco-resistance (*gabrb3*) can be recapitulated in zebrafish models.

Although, CRISPR/Cas9 works with remarkable efficiency to disrupt gene function in zebrafish^39,40^, recent large-scale efforts have not reported on epilepsy or clinically-relevant functional outcome measures^16,17^. To present robust and well-controlled functional assays, we outcrossed all EZP lines a minimum of three generations and blindly analyzed homozygous, heterozygous and WT siblings. This approach avoids off-target or toxicity effects from microinjection or CRISPR/Cas9 editing that might cause identification of false positives. A limitation typical of these types of CRISPR-based larval zebrafish studies, focused primarily on novel genes, is that the full spectrum of tools (antibodies, etc.) or functional assays (single-cell electrophysiology) necessary to confirm LOF mutation are not available. Nonetheless, epileptic activities seen in CRISPR/Cas9 deficient (*aldh7a1*)^46,55^ or ENU-generated (*scn1lab^s552/s552^*)^23^ zebrafish were successfully recapitulated here. Interestingly, but perhaps not surprisingly, the majority of our CRISPR-generated single gene LOF zebrafish mutants were not associated with epilepsy phenotypes at this stage of larval development (5-6 dpf). It is possible that many of these single gene mutations are one factor in the emergence of epilepsy in humans, but full clinical phenotypes rely upon polygenic factors^56,57^, epigenetics^53^ or environmental issues such as early-life febrile seizures^58^. Developmental considerations are an additional confounding factor^2,3^, as clear epileptic phenotypes may emerge at later juvenile or adult timepoints. Although a potential limitation for interpretation of these studies, we chose to focus this initial phenotypic screening effort on larval developmental ages that would lend themselves to future high-throughput drug discovery. Where single gene mutant mice are available for electrophysiology comparisons a similar lack of unprovoked seizure phenotypes have been reported e.g., *Cdkl5^59,60^*, *Chd2^61^* or *Depdc5^62^*. Further, the frequency and severity of seizure activity in patients with single gene mutations can also be variable e.g., *SCN8^63^, PCDH19^64^*, *MEF2C^65^*, *CDKL5* and *ARX^66^*, which highlights the complexity of modelling rare epilepsy gene candidates.

Our previous studies established the presence of hyperactive and seizure-like (stage III) behaviors in PTZ-treated WT larvae and spontaneously in *scn1lab^s552/s552^* mutant larvae, a model of Dravet syndrome^23,44^. These stage III behaviors are defined as brief clonus-like convulsions followed by a loss of posture, where a larvae falls on its side and remains immobile for 1–3 s (manually scored)^44^. Behavioral readouts were instrumental in primary screens aimed at finding novel anti-epileptic drugs that treat Dravet syndrome, ultimately allowing us to test over 3500 drugs in less than 5 years^23,24,27^ and advancing our lead candidate to clinical trials (https://clinicaltrials.gov/ct2/show/NCT04462770). Here, we further refine our definition of seizure-like movements as events ≥ 28 mm/s in velocity and ≥1 s in duration and created a MATLAB algorithm to efficiently detect these events in our behavioral assays; total distance moved was not a reliable measure of these events. Interestingly, of our 8 EZP-epilepsy CRISPR lines, only the most robust phenotypic line (*scn1lab* mutants) had significantly more seizure-like behavioral events compared to controls, suggesting that hyper-locomotion alone may not be sufficient to identify epileptic phenotypes. Interestingly, hypo-locomotion seen here in *arxa* mutant larvae [also reported in *tsc2^67^* and *gabrg2^68^* mutants, respectively] may represent a pathological behavioral state. Ultimately and mimicking clinical diagnoses of epilepsies using video-electroencephalographic monitoring^2,30,44^, our electrophysiology-based screening approach successfully identified epileptic activity that was not easily detected in locomotion-based assays. Although simple locomotor readouts have grown popular as seizure assays^69–74^, this study emphasizes the rigor necessary to accurately identify epileptic phenotypes in zebrafish and suggests that sole reliance on behavior may lead to misleading conclusions during phenotyping and/or drug discovery efforts.

Overall, the Epilepsy Zebrafish Project demonstrates the power of large-scale phenotype-based analyses of human gene mutations and all mutant lines are available to the scientific community (https://zebrafishproject.ucsf.edu). These CRISPR-generated zebrafish models have two important advantages: first, they provide a valuable *in vivo* model system to explore underlying pathophysiological mechanisms in rare genetic epilepsies. Second, they provide an easily accessible preclinical model system for high-throughput drug discovery and therapy development that is far more efficient than rodent models. Pilot neurodevelopmental and pharmacological data was provided for several epileptic zebrafish lines here as a potential starting point for further investigations. We anticipate, and hope, that future studies using these zebrafish will help us to better understand genetic disorders and further the ultimate vision of precision medicine.

## Methods

### Zebrafish Husbandry

All procedures described herein were performed in accordance with the Guide for the Care and Use of Animals (ebrary Inc., 2011) and adhered to guidelines approved by the University of California, San Francisco Institution Animal Care and Use Committee (IACUC approval #: AN171512-03A). The zebrafish lines were maintained in a temperature-controlled facility on a 14:10 hour light:dark cycle (9:00 AM −11:00 PM PST). Juvenile and adult zebrafish were housed on aquatic units with an automated feedback control unit that maintained the system water conditions within the following ranges: temperature; 28-30 °C, pH; 7.5-8.0 and conductivity; 690-740 mS/cm. Juveniles (30-60 dpf) were fed twice daily, once with JBL powder (JBL NovoTom Artemia) and the other with JBL powder + live brine shrimp (Argent Aquaculture). Older juveniles and adults were also fed two times per day, first with flake food (tropical flakes, Tetramin) and then with flake food and live brine shrimp. Zebrafish embryos and larvae were raised in an incubator kept at 28.5 °C under the same light-dark cycle as the facility. The solution or ‘embryo medium’ used for the embryos and larvae consisted of 0.03% Instant Ocean (Aquarium Systems, Inc.) and 0.000002% methylene blue in reverse osmosis-distilled water. Larvae were fed with powder (6-10 dpf) or JBL powder + brine shrimp (11-29 dpf).

### Zebrafish homology prediction

To improve our confidence in modeling epilepsy at the genetic level in zebrafish, we established a zebrafish homology score. To determine the homology score the percent protein identity and DIOPT score was used. The percent protein identity was established from Ensembl (GRCz10) using the predicted human orthologue gene. When the human orthologue gene was not predicted by Ensembl, a Clustal Omega analysis was performed using standard parameters. The DIOPT score was established using the MARRVEL (http://marrvel.org/) database and is the number of orthologue prediction tools that predicted a given orthologue pair. Twelve orthologue prediction tools (Comara, Eggnog, Homologene, Inparanoid, OMA, OrthoDB, orthoMCL, Panther, Phylome, RoundUP, TreeFam and ZFIN) were used to predict zebrafish orthologs. The homology score represents the average of the percent identity and the DICOT score as a percentage. A gene with a homology score >65 was considered for the EZP.

### Zebrafish gene expression analysis

Adult tissue expression was determined using the Phylofish database^75^. Development expression was determined using semi-quantitative RT-PCR. Pools of 25 to 50 zebrafish embryos or larvae were collected at 4-cell, 32-cell, high, sphere, 12 hpf, 1, 2, 3, 4, 5, and 7 dpf for expression analysis. Total mRNA was extracted from whole embryos or larvae using a phenol/chloroform extraction protocol. After extraction, 1 μg of purified RNA was treated with DNaseI and retrotranscribed to cDNA using following SuperScript IV Reverse Transcriptase (8091050, Invitrogen) the manufacturer's protocol. The temporal expression of genes was characterized RT-PCR using GoTaq Master Mix (M712C, Promega) and oligonucleotide sequences are listed at https://zebrafishproject.ucsf.edu. Thermal cycling conditions included an initial denaturation at 95°C for 5 min, followed by 40 cycles at 95°C for 30 sec, 56°C for 30 sec, and 72°C for 30 sec and a final incubation at 72°C for 7 min.

### Generation of CRISPR mutant lines

Zebrafish mutant lines of the 40 genes were generated using CRISPR-Cas gene editing in Tupfel Long-Fin (TL) wild-type zebrafish (ZIRC). CRISPRScan was used to identify sgRNA sequences with high predicted cut efficiencies for early exons and sgRNAs were synthesized using T7 in vitro transcription with the MEGAshortscriptTM T7 Transcription Kit (AM1354, ThermoFisher). To minimize off target-effects, we selected target sites with the lowest number of potential mutagenesis and with a minimum of three mismatches with every other site in the genome. Fertilized embryos (1-2 cell stage) were co-injected with ~2 nl of sequence-specific sgRNA (~10-25 ng/μl), Cas9 mRNA (~250 ng/μl) and 0.4% rhodamine b. At 1 dpf, embryos were sorted for fluorescence and genomic DNA extracted using Zebrafish Quick Genotyping DNA Preparation Kit (GT02-02, Bioland Scientific) from pools of 5-10 healthy, microinjected and un-injected larvae. The samples were Sanger sequenced to assess gene editing at the guide target site. Once editing was confirmed, the remaining embryos were raised to adulthood. Resulting F0 mosaic adults, confirmed by Sanger sequencing DNA from fin-clips, were crossed with TL zebrafish to create stable heterozygote F2 and greater generations of breeders for our experiments. Guide RNA, primer sequences and PCR protocols for all lines can be found in Supplementary Table 3. All experiments were done blinded using unfed larvae between 3-14 dpf. At this stage larvae are sexually indistinguishable.

### Electrophysiology

Zebrafish larvae (5-6 dpf) were randomly selected, briefly exposed to cold anesthesia or pancuronium (300 μM) and immobilized, dorsal side up, in 2% low-melting point agarose (BP1360-100, Fisher Scientific) within a vertical slice perfusion chamber (Siskiyou Corporation, #PC-V). Slice chambers containing one or two larvae, were placed on the stage of an upright microscope (Olympus BX-51W) and monitored continuously using a Zeiss Axiocam digital camera. Under visual guidance, gap-free local field potential recordings (LFP; 15 min duration) were obtained from optic tectum using a single glass microelectrode (*WPI glass #TW150 F-3*); ~ 1 μm tip diameter; 2 mM NaCl internal solution), as described^43,44^. LFP voltage signals were low-pass filtered at 1 kHz (−3 dB; eight-pole Bessel), digitized at 10 kHz using a Digidata 1320 A/D interface (Molecular Devices) and stored on a PC computer running AxoScope 10.3 software (Molecular Devices). For pharmacology experiments, continuous gap-free LFP recordings were made for 1 hr and drug concentrations are based on previously published data^23,44^. Larvae were gently freed from agarose at the conclusion of recording epochs for *post hoc* genotyping by investigators blind to status of the experiment. Electrophysiology files were also coded for *post hoc* analysis off-line. Experiments were performed on at least three independent clutches of larvae for each line; a minimum of 75 larvae were screened per line. Individual abnormal electrographic seizure-like events were defined as: (i) brief interictal-like events comprised of spike upward or downward membrane deflections greater than 3x baseline noise level or (ii) long duration, large amplitude ictal-like multi or poly-spike events greater than 5x baseline noise level. Quantification of epileptiform events was performed using Clampfit 10.3 (Molecular Devices) or custom MATLAB (MathWorks; Figure 3) software by investigators blind to status of the experiment. A binning method combined with a sliding window algorithm was used to calculate the active level of the signal within the current time window. The value of each bin was used to identify the start and end of an event. We used a range of voltage thresholds (0.15 – 0.25 mV, depending on the noise level) and a relative threshold (3x Standard Deviation) for detection of interictal events. By comparing manual-auto counting results of a testing data sample for each recording (Figure 4d), we fine-tuned the threshold detection for each recording to a level where auto counting results were close to the manual counting results (< 3% difference). All files were un-coded and combined with genotyping data at the end of this process.

### Larval Behavior

#### Basal locomotion

Behavioral studies conducted on select EZP lines utilized a 96-well format and automated locomotion detection using a DanioVision system running EthoVision XT 11.5 software (DanioVision, Noldus Information Technology). Zebrafish larvae were transferred from their home incubator to the test room at least 10 min before the experiment. After larvae were individually transferred to wells in ~150 μl of embryo media, the 96 well-plate was placed in the DanioVision observation chamber and left undisturbed for 1 hr. Larval movement was tracked for 15 min at 25 frames per sec with the following detection settings: method; DanioVision, sensitivity; 110, video pixel smoothing; low, track noise reduction; on, subject contour; 1 pixel (contour dilation, erode first then dilate), subject size; 4-4065. For each zebrafish line, experiments were performed with at least 3 different clutches and *post hoc* genotyping. Mean and maximum velocity of each larvae were calculated. Additionally, high-speed seizure behaviors were scored using a MATLAB algorithm developed by our laboratory and validated on PTZ and *scn1lab* seizure models (Figure 7).

#### Seizure-like behavioral event classification

To classify larval movements, we first processed the videos with EthoVision software 11.5 (Noldus) to identify a larva’s position at an acquisition rate of 25 frames/sec, using the same detection settings listed in the ‘basal locomotion’ assay, except with the track noise reduction off. Using custom-written MATLAB-based software, we then extracted movement events defined as times when larvae speed exceeded a threshold of 0.9 mm/sec for at least 160 msec. Adjacent events were combined if the time interval was less than 40 msec. Furthermore, when the maximum speed within an event was lower or higher than a cutoff threshold, the movement events were classified into low- and high- speed events, respectively. For the analysis in Figure 7, we calculated the distribution of all movements in a large control group of larvae and then identified the speed value threshold at 1.5x Standard Deviation to be used as a cutoff threshold, unless otherwise specified. Similar results for larval WT movement speeds and duration have been previously reported^76^. Seizure-like events were defined as high-speed movement events that lasted longer than 1 sec validated on PTZ and *scn1lab* seizure models.

#### Behavioral effects of electrode implantation

WT larvae (5 dpf) in 100 mm petri dishes were transferred to the test room and subjected to one of three treatments:

<u>Treatment 1</u>: Larvae were briefly anesthetized in pancuronium (300 μM) and then immobilized in 2% agarose dorsal side up on a recording chamber. About 3 ml of recording media was added to the chamber then a glass micro-electrode was positioned in the forebrain for LFP recording as previously described^43,44^. After 15-30 min, the electrode was removed and the larva gently released from agarose and transferred to a petri dish with embryo medium.
<u>Treatment 2</u>: Larvae were briefly anesthetized in pancuronium (300 μM) and then immobilized in 2% agarose dorsal side up on a recording chamber. After 15-30 min, the larvae were gently released from the agarose and transferred to a petri dish with embryo medium.
<u>Treatment 3</u>: Larvae were left undisturbed in original petri dish.

At the end of the experiment, all treatment groups were returned to the home incubator until behavioral experiments. Four hours after treatment, larvae were returned to test room and left undisturbed for 10 min. Larvae were individually transferred to a 96 well plate in ~150 μl of embryo media and the plate then placed in the DanioVision observation chamber. After 15 min, larval movement was tracked for 30 min using settings outlined in ‘basal locomotion’. Once completed, the plate was removed and returned to the home incubator. The same steps were followed to record behavior 24 hr post-treatment.

### Survival Assay

For each line, 20-24 zebrafish larvae were randomly selected from at least two clutches and were placed in a 100 mm petri dish containing ~40 ml egg water. The larvae were monitored twice daily and dead larvae were lysed using Bioland Zebrafish Quick Lysis Kit. Larvae were not fed throughout the duration of the assay. This was done to eliminate potential effects of variations in larval feeding, ultimately providing us with a robust method to identify early-stage larval lethality phenotypes. Unfed larvae typically die by 12 dpf^77^. Samples were genotyped using protocols specified in Supplementary Table 3.

#### Pyridoxine supplementation: aldh7a1 survival

At 4 dpf, larvae were placed individually in 24 well plate with 500 μl 10 mM pyrodixine or egg water (control). Treatment was removed and replaced with fresh egg water. Larvae were then treated with 500 μl 10 mM pyrodixine or egg water (control) for 30 min daily. During daily monitoring, dead larvae were lysed using Bioland Zebrafish Quick Lysis Kit. Samples were genotyped using protocols specified in Supplementary Table 3.

### Imaging

For morphology measurements in the *eef1a2* CRISPR line, larvae were placed individually in one well of a μ-well microscope slide (iBidi) and high-resolution images obtained using an optiMOS CMOS camera (QImaging) camera mounted on a SteREO Discovery.V8 stereomicroscope (Zeiss). Files were coded and processed by an investigator blind to status of the experiment. Larvae were collected for independent *post hoc* genotyping at the conclusion of image acquisition. Images were analyzed by a third investigator using DanioScope software (Noldus, version 1.0.109). Standard head (overall head length, midbrain and forebrain widths) and body length (distance from anterior tip of head to base of caudal fin) measurements were obtained. Files were un-coded and combined with genotyping data at the end of this process.

#### Interneuron Quantification

For imaging studies, *arxa* CRISPR line was crossed with a *dlx5a-dlx6a:GFP:nacre* transgenic zebrafish line provided by Marc Ekker^52^. For analysis of interneuron density in *arxa* WT and homozygote mutants, green fluorescent protein (GFP)-expressing larvae were sorted by fluorescence at 2 to 3 dpf and imaged at 5 dpf using a Zeiss Z.1 light sheet microscope with a 20X objective. Zebrafish were anesthetized in 0.04% tricaine mesylate and embedded in 2% low melting point agarose inside a glass capillary. The imaging sample chamber was filled with embryo medium. Z-stack images were acquired at 5 μm intervals starting at the first visible dorsal GFP-positive cell. Following image acquisition, larvae were gently removed from agar and independently genotyped. Imaging files were coded and analyzed *post hoc* by an investigator blind to status of the experiment. Images were then processed in Fiji (ImageJ)^78^. Neurons were quantified with an algorithm modified from “3D watershed technique” (ImageJ macro developed by [Bindokas V, 17-September-2014. Available: https://digital.bsd.uchicago.edu/%5Cimagej_macros.html.]).

### Statistical analysis

Statistical tests were performed using MATLAB or GraphPad Prism. One-way ANOVA with Dunnett’s multiple comparison tests or non-parametric *t* tests were used. Data are presented as mean ± S.E.M. Individual analyses are described in Results.

### Data and software availability

All custom MATLAB programs will be made available upon reasonable request. Representative electrophysiology tracings, Kaplan-Meier survival plots, behavioral data, sequencing information are available on our web-portal (https://zebrafishproject.ucsf.edu). The datasets generated during the current studies are available from the corresponding author on reasonable request.

## Reporting summary

Further information on research design is available in the Nature Research Reporting Summary linked to this article.

## Acknowledgements

We would like to thank Dan Lowenstein, Gemma Carvill and Robert Hunt for comments and feedback on conceptualization of the Epilepsy Zebrafish Project. We thank Kathryn Salvati and Mark Beenhakker for sharing MATLAB code for generation of time-frequency histograms. We thank Sarai Diaz and Ifechukwu Okeke for assistance on maintenance of zebrafish lines. This work was supported by NIH/NINDS R01 award #NS103139 (to S.C.B.); International Foundation for CDKL5 Research and Bow Foundation grants (to S.C.B); Lennox-Gastaut Syndrome Foundation fellowships (to B.G. and C.C); and a Dravet Syndrome Foundation fellowship (to A.G.).

## Author Contributions

A.G. & S.C.B. conceived the project. A.G. & C.C. designed the CRISPR strategy and trained subsequent personnel on generation of mutant lines. A.G. & C.C. directed and supervised the research. A.G., C.C., J.L., B.G., & K.H. generated mutant zebrafish lines. A.G., K.H. & M.A. conducted the gene expression studies. C.C. designed and guided the behavioral assay studies. C.C. & C.O. collected the behavioral data and C.C., C.O. & R. P. analyzed the data. S.C.B., M.M., & M.T.D. collected and analyzed electrophysiology data. T.Q. & J.L. collected and analyzed light sheet microscopy data. C.C., M.M., & F.F. collected and analyzed survival data. J.L. & R.P. designed and wrote MATLAB programs for data analysis. M.M. & C.O. created the database website. M.T.D., M.A., C.O. & F.F. maintained the zebrafish colony. A.G., C.C. & S.C.B. wrote and edited the paper.

## Additional information

Supplementary Information is available for this paper.

---

**Supplementary Figure 1.**
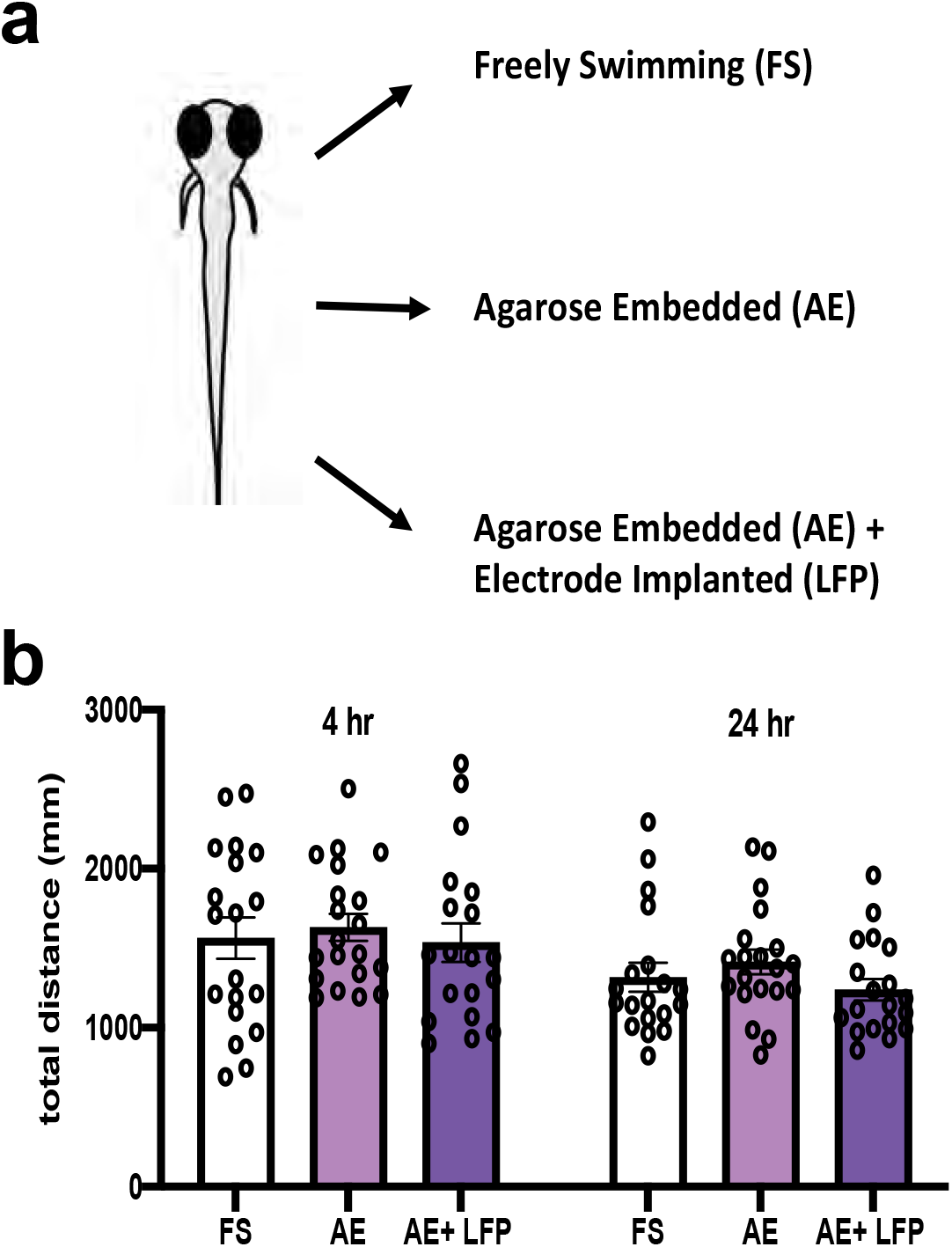
Local field potential recordings are minimally invasive. (**a**) Five dpf larvae were left freely swimming in embryo medium or subjected to agar embedding or agar embedding with electrode implantation, and behavior was tracked 4 hr and 24 hr after each treatment. Results show no significant differences in the total distance traveled (**b**) or maximum velocity (data not shown) of larvae when compared across the treatment groups. Data displayed as mean ± SEM.

**Supplementary Figure 2.**
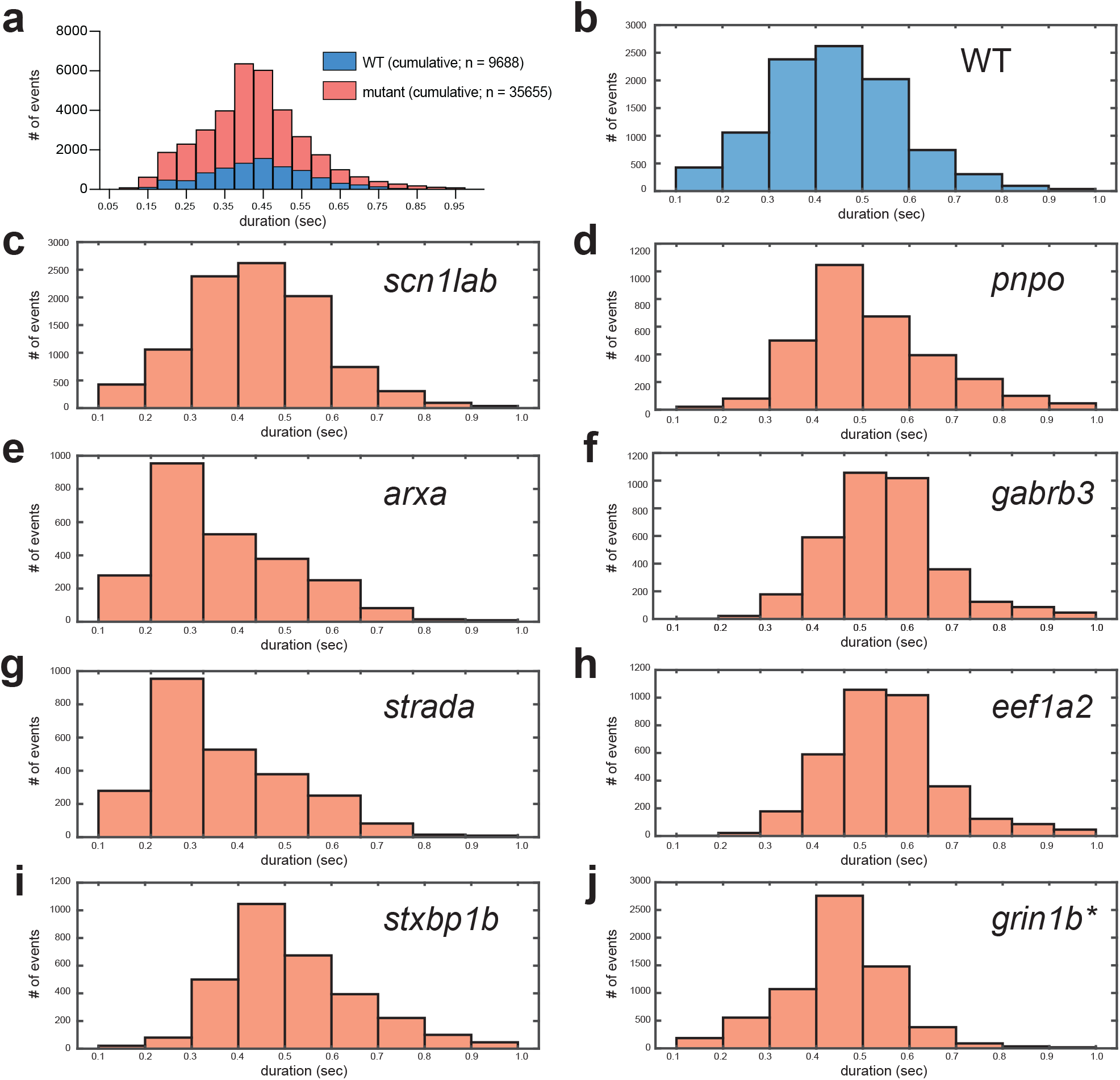
Distribution of ictal events. Histograms depict number and duration of ictal events cumulatively (**a-b**) and across individual EZP-epilepsy lines (**c-j**). Asterisk for *grin1b* designates heterozygote. Interictal events were measured using a custom MATLAB-based program for EZP-epilepsy lines and WT siblings.

**Supplementary Figure 3.**
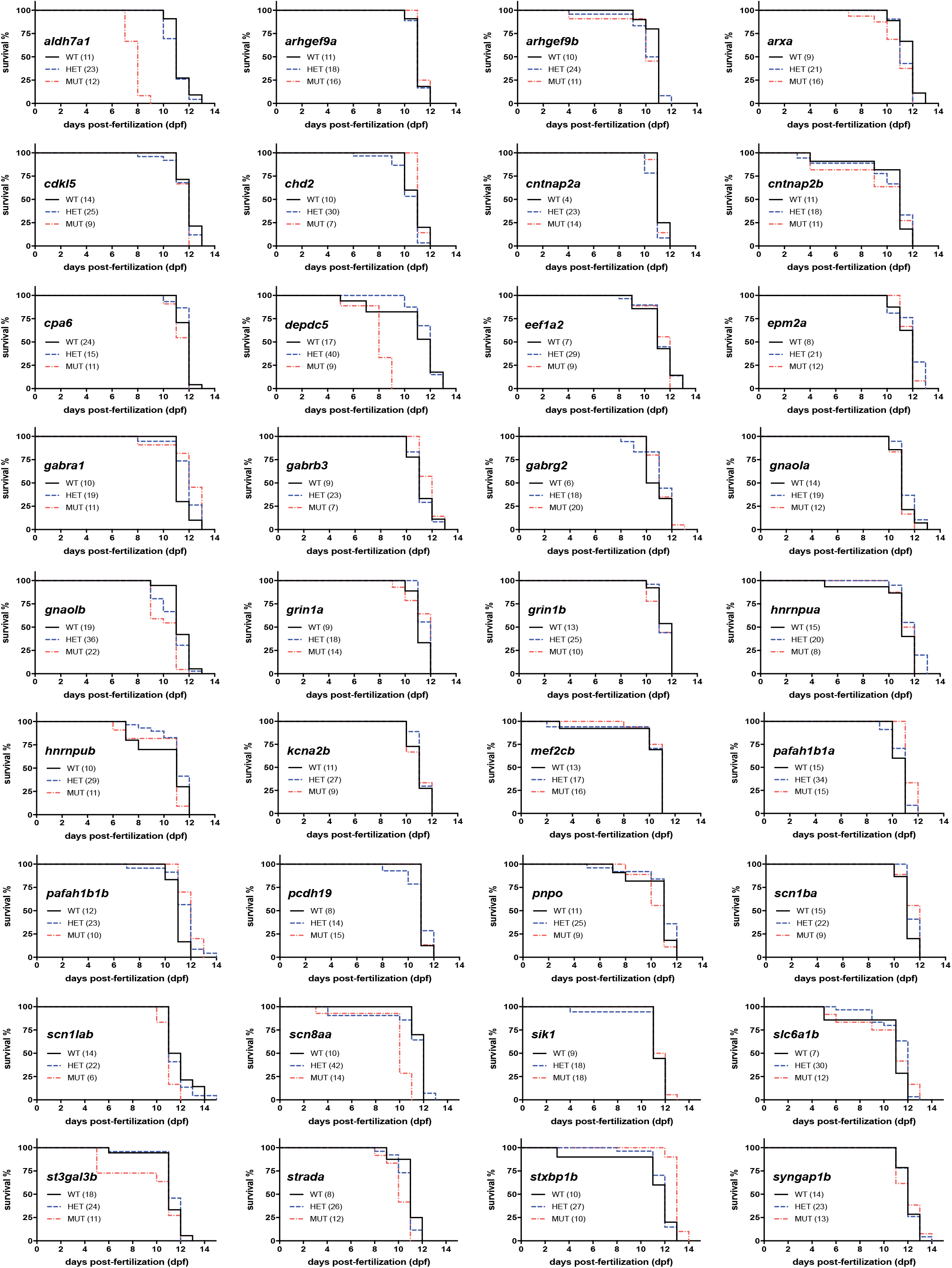
Kaplan-Meier survival curves for zebrafish CRISPR lines. Plots of survival for unfed WT, heterozygous and homozygous larvae across all zebrafish lines.

**Supplementary Figure 4.**
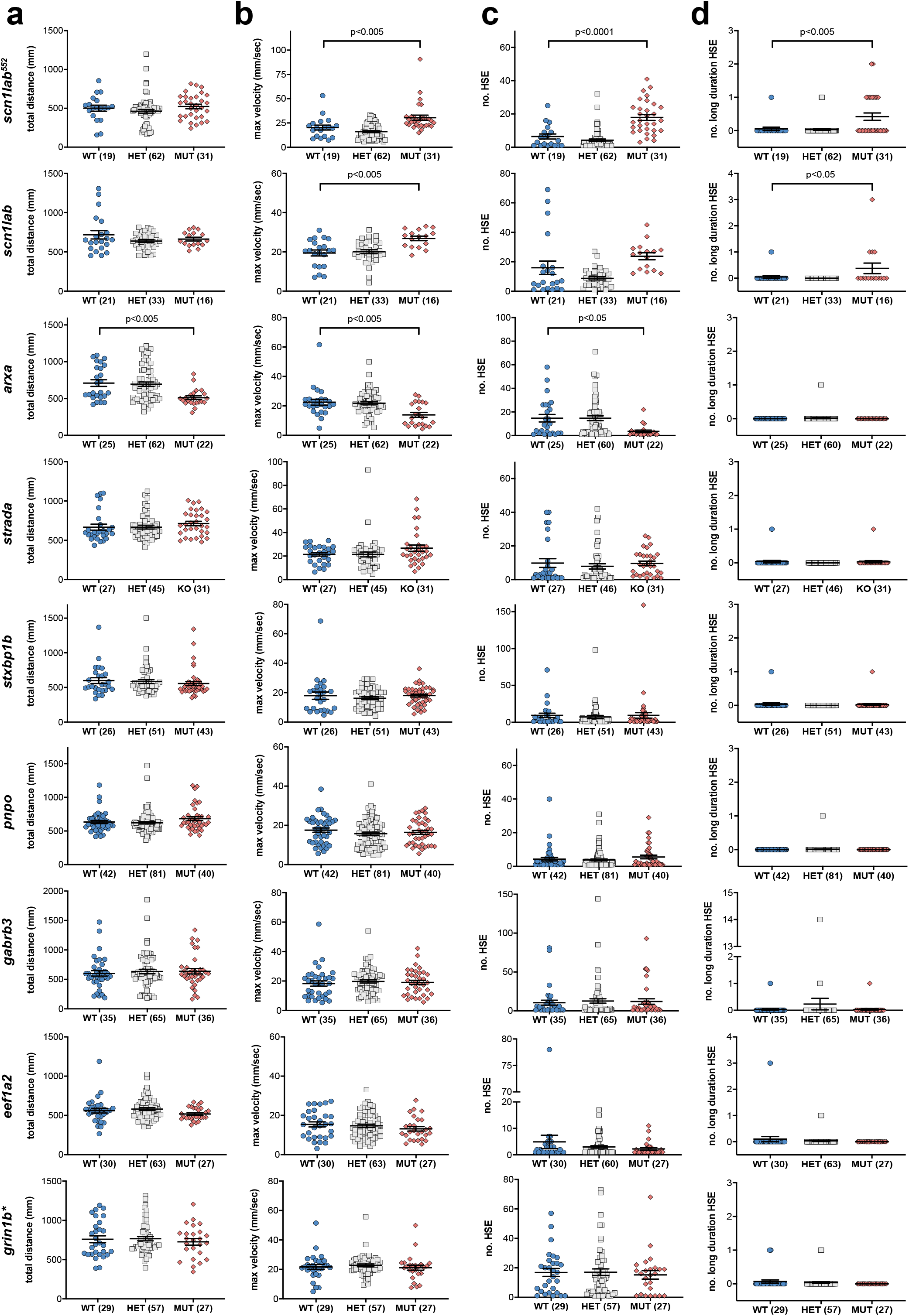
Basal locomotor activity of epileptic zebrafish lines. Five dpf larval zebrafish were tracked in the behavioral assay and graphs depict (**a**) total distance traveled, (**b)** maximum velocity, (**c**) number of high-speed events (HSE) and (**d**) number of long duration HSE observed across the various lines. Total distance and maximum velocity were extracted directly from EthoVision XT 11.5 software while the number of events ≥ 28 mm/s (HSE) and long duration HSE (≥1 s) were scored using an in-house MATLAB algorithm. Data displayed as scatter plots showing individual larval values and error bars represent mean and SEM. Statistics calculated using One-way ANOVA and *post hoc* Dunnett multiple comparison test, \**p* ≤ 0.05, \*\**p*≤ 0.005, \*\**p* < 0.0001.

**Supplementary Figure 5.**
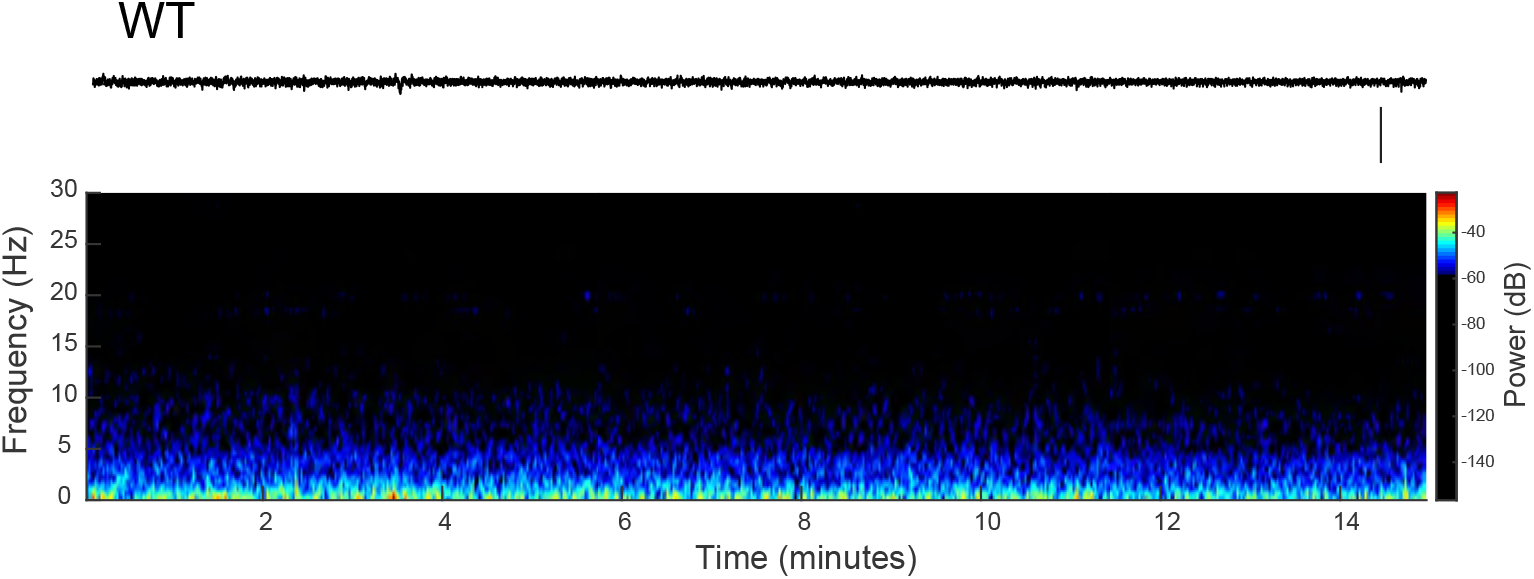
Wild-type recording. Representative raw LFP recording trace (top) along with a corresponding wavelet time-frequency spectrogram (bottom) for a representative WT zebrafish larvae. Scale bar = 500 μV.

**Table S1:**
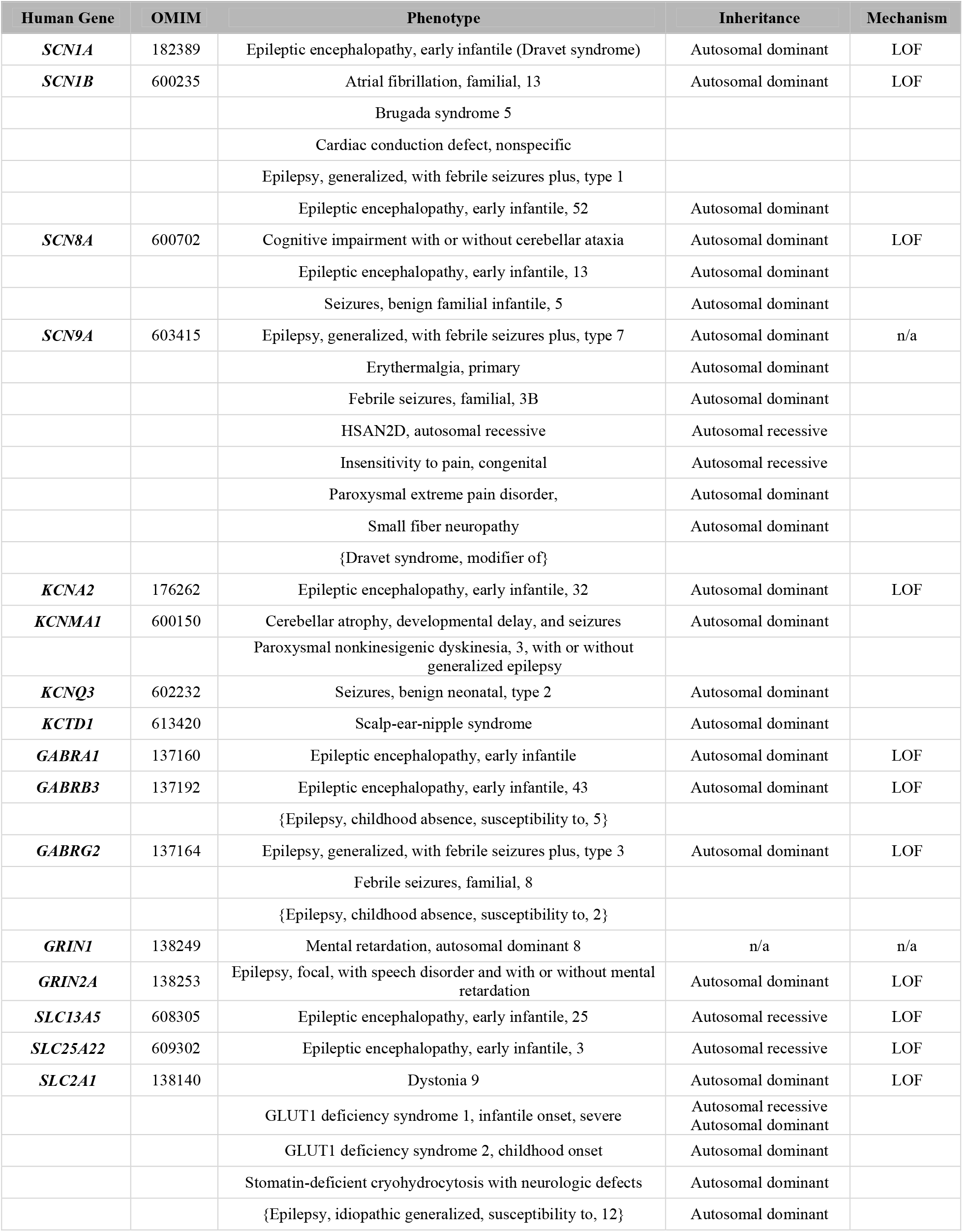
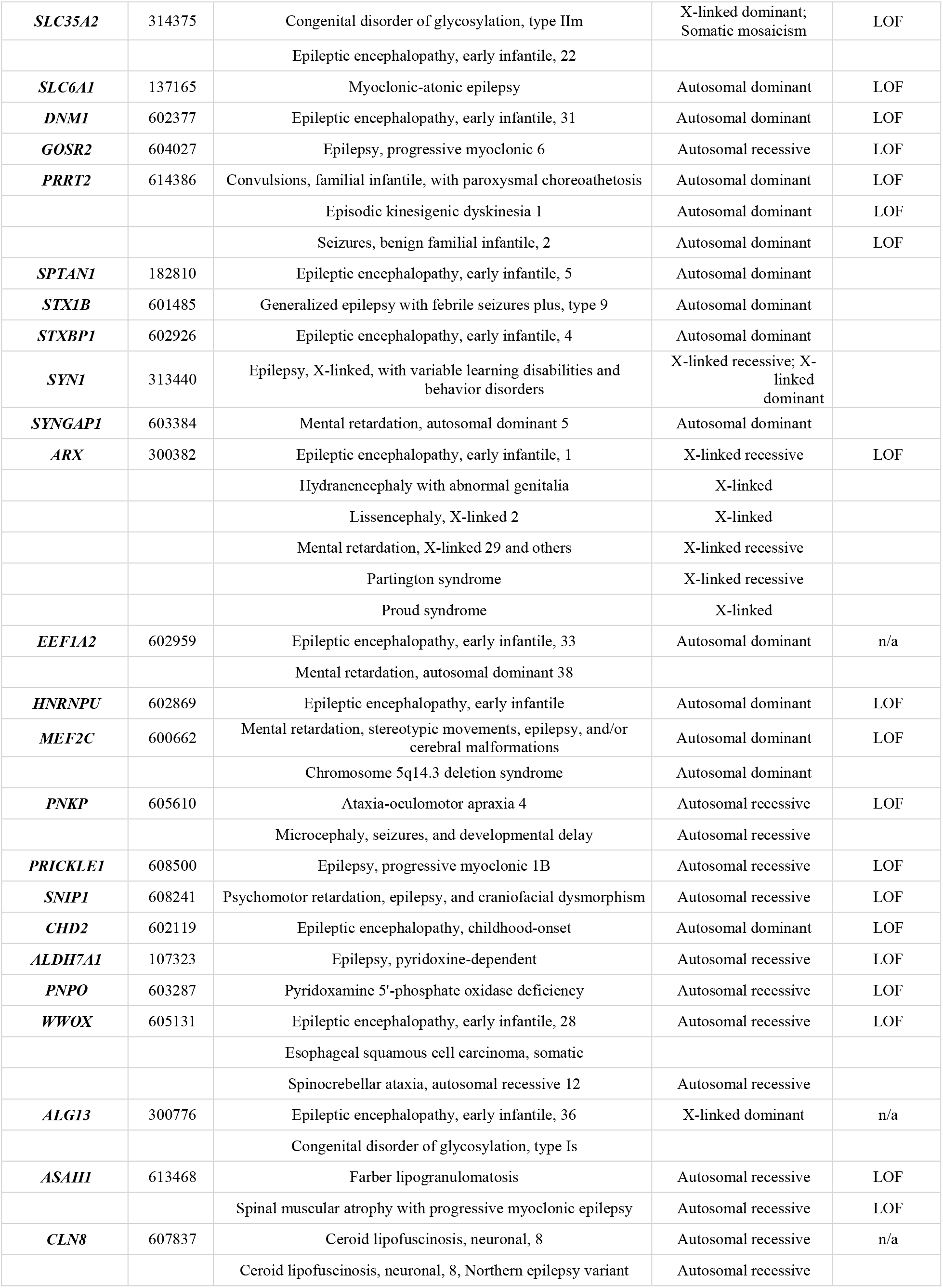
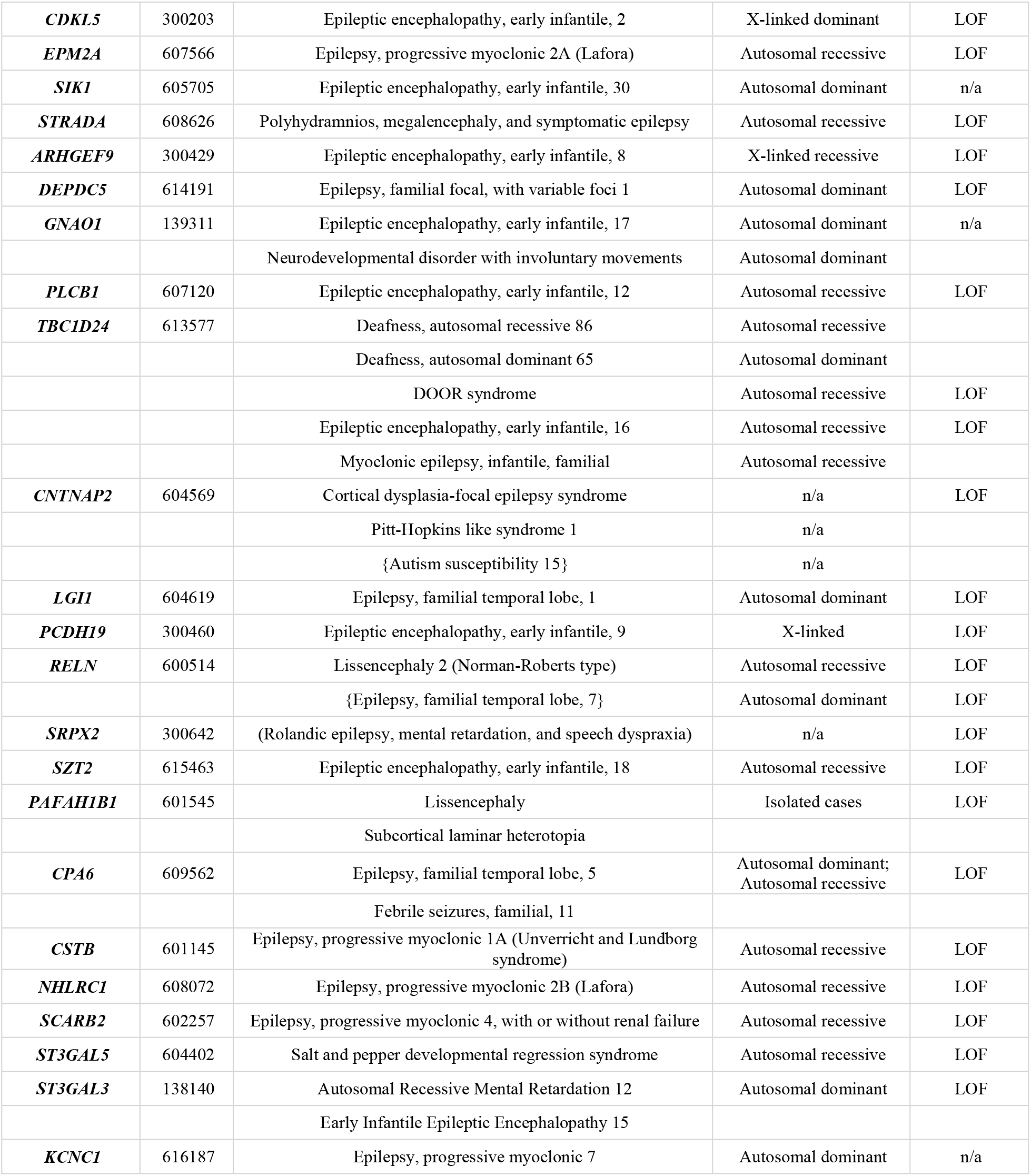
Genes associated with an epilepsy phenotype that were considered for the Epilepsy Zebrafish Project.

| Human Gene | OMIM | Phenotype | Inheritance | Mechanism |
| --- | --- | --- | --- | --- |
| *SCN1A* | 182389 | Epileptic encephalopathy, early infantile (Dravet syndrome) | Autosomal dominant | LOF |
| *SCN1B* | 600235 | Atrial fibrillation, familial, 13 | Autosomal dominant | LOF |
|  |  | Brugada syndrome 5 |  |  |
|  |  | Cardiac conduction defect, nonspecific |  |  |
|  |  | Epilepsy, generalized, with febrile seizures plus, type 1 |  |  |
|  |  | Epileptic encephalopathy, early infantile, 52 | Autosomal dominant |  |
| *SCN8A* | 600702 | Cognitive impairment with or without cerebellar ataxia | Autosomal dominant | LOF |
|  |  | Epileptic encephalopathy, early infantile, 13 | Autosomal dominant |  |
|  |  | Seizures, benign familial infantile, 5 | Autosomal dominant |  |
| *SCN9A* | 603415 | Epilepsy, generalized, with febrile seizures plus, type 7 | Autosomal dominant | n/a |
|  |  | Erythralgia, primary | Autosomal dominant |  |
|  |  | Febrile seizures, familial, 3B | Autosomal dominant |  |
|  |  | HSAN2D, autosomal recessive | Autosomal recessive |  |
|  |  | Insensitivity to pain, congenital | Autosomal recessive |  |
|  |  | Paroxysmal extreme pain disorder, | Autosomal dominant |  |
|  |  | Small fiber neuropathy | Autosomal dominant |  |
|  |  | {Dravet syndrome, modifier of} |  |  |
| *KCNA2* | 176262 | Epileptic encephalopathy, early infantile, 32 | Autosomal dominant | LOF |
| *KCNMA1* | 600150 | Cerebellar atrophy, developmental delay, and seizures | Autosomal dominant |  |
|  |  | Paroxysmal nonkinesigenic dyskinesia, 3, with or without generalized epilepsy |  |  |
| *KCNQ3* | 602232 | Seizures, benign neonatal, type 2 | Autosomal dominant |  |
| *KCTD1* | 613420 | Scalp-ear-nipple syndrome | Autosomal dominant |  |
| *GABRA1* | 137160 | Epileptic encephalopathy, early infantile | Autosomal dominant | LOF |
| *GABRB3* | 137192 | Epileptic encephalopathy, early infantile, 43 | Autosomal dominant | LOF |
|  |  | {Epilepsy, childhood absence, susceptibility to, 5} |  |  |
| *GABRG2* | 137164 | Epilepsy, generalized, with febrile seizures plus, type 3 | Autosomal dominant | LOF |
|  |  | Febrile seizures, familial, 8 |  |  |
|  |  | {Epilepsy, childhood absence, susceptibility to, 2} |  |  |
| *GRIN1* | 138249 | Mental retardation, autosomal dominant 8 | n/a | n/a |
| *GRIN2A* | 138253 | Epilepsy, focal, with speech disorder and with or without mental retardation | Autosomal dominant | LOF |
| *SLC13A5* | 608305 | Epileptic encephalopathy, early infantile, 25 | Autosomal recessive | LOF |
| *SLC25A22* | 609302 | Epileptic encephalopathy, early infantile, 3 | Autosomal recessive | LOF |
| *SLC2A1* | 138140 | Dystonia 9 | Autosomal dominant | LOF |
|  |  | GLUT1 deficiency syndrome 1, infantile onset, severe | Autosomal recessive<br>Autosomal dominant |  |
|  |  | GLUT1 deficiency syndrome 2, childhood onset | Autosomal dominant |  |
|  |  | Stomatin-deficient cryohydrocytosis with neurologic defects | Autosomal dominant |  |
|  |  | {Epilepsy, idiopathic generalized, susceptibility to, 12} | Autosomal dominant |  |
| **SLC35A2** | 314375 | Congenital disorder of glycosylation, type IIIm | X-linked dominant;<br>Somatic mosaicism | LOF |
|  |  | Epileptic encephalopathy, early infantile, 22 |  |  |
| **SLC6A1** | 137165 | Myoclonic-atonic epilepsy | Autosomal dominant | LOF |
| **DNM1** | 602377 | Epileptic encephalopathy, early infantile, 31 | Autosomal dominant | LOF |
| **GOSR2** | 604027 | Epilepsy, progressive myoclonic 6 | Autosomal recessive | LOF |
| **PRRT2** | 614386 | Convulsions, familial infantile, with paroxysmal choreoathetosis | Autosomal dominant | LOF |
|  |  | Episodic kinesigenic dyskinesia 1 | Autosomal dominant | LOF |
|  |  | Seizures, benign familial infantile, 2 | Autosomal dominant | LOF |
| **SPTAN1** | 182810 | Epileptic encephalopathy, early infantile, 5 | Autosomal dominant |  |
| **STX1B** | 601485 | Generalized epilepsy with febrile seizures plus, type 9 | Autosomal dominant |  |
| **STXBP1** | 602926 | Epileptic encephalopathy, early infantile, 4 | Autosomal dominant |  |
| **SYN1** | 313440 | Epilepsy, X-linked, with variable learning disabilities and behavior disorders | X-linked recessive; X-linked dominant |  |
| **SYNGAP1** | 603384 | Mental retardation, autosomal dominant 5 | Autosomal dominant |  |
| **ARX** | 300382 | Epileptic encephalopathy, early infantile, 1 | X-linked recessive | LOF |
|  |  | Hydranencephaly with abnormal genitalia | X-linked |  |
|  |  | Lissencephaly, X-linked 2 | X-linked |  |
|  |  | Mental retardation, X-linked 29 and others | X-linked recessive |  |
|  |  | Partington syndrome | X-linked recessive |  |
|  |  | Proud syndrome | X-linked |  |
| **EEF1A2** | 602959 | Epileptic encephalopathy, early infantile, 33 | Autosomal dominant | n/a |
|  |  | Mental retardation, autosomal dominant 38 |  |  |
| **HNRNPU** | 602869 | Epileptic encephalopathy, early infantile | Autosomal dominant | LOF |
| **MEF2C** | 600662 | Mental retardation, stereotypic movements, epilepsy, and/or cerebral malformations | Autosomal dominant | LOF |
|  |  | Chromosome 5q14.3 deletion syndrome | Autosomal dominant |  |
| **PNKP** | 605610 | Ataxia-oculomotor apraxia 4 | Autosomal recessive | LOF |
|  |  | Microcephaly, seizures, and developmental delay | Autosomal recessive |  |
| **PRICKLE1** | 608500 | Epilepsy, progressive myoclonic 1B | Autosomal recessive | LOF |
| **SNIP1** | 608241 | Psychomotor retardation, epilepsy, and craniofacial dysmorphism | Autosomal recessive | LOF |
| **CHD2** | 602119 | Epileptic encephalopathy, childhood-onset | Autosomal dominant | LOF |
| **ALDH7A1** | 107323 | Epilepsy, pyridoxine-dependent | Autosomal recessive | LOF |
| **PNPO** | 603287 | Pyridoxamine 5'-phosphate oxidase deficiency | Autosomal recessive | LOF |
| **WWOX** | 605131 | Epileptic encephalopathy, early infantile, 28 | Autosomal recessive | LOF |
|  |  | Esophageal squamous cell carcinoma, somatic |  |  |
|  |  | Spinocerebellar ataxia, autosomal recessive 12 | Autosomal recessive |  |
| **ALG13** | 300776 | Epileptic encephalopathy, early infantile, 36 | X-linked dominant | n/a |
|  |  | Congenital disorder of glycosylation, type Is |  |  |
| **ASAH1** | 613468 | Farber lipogranulomatosis | Autosomal recessive | LOF |
|  |  | Spinal muscular atrophy with progressive myoclonic epilepsy | Autosomal recessive | LOF |
| **CLN8** | 607837 | Ceroid lipofuscinosis, neuronal, 8 | Autosomal recessive | n/a |
|  |  | Ceroid lipofuscinosis, neuronal, 8, Northern epilepsy variant | Autosomal recessive |  |
| **CDKL5** | 300203 | Epileptic encephalopathy, early infantile, 2 | X-linked dominant | LOF |
| **EPM2A** | 607566 | Epilepsy, progressive myoclonic 2A (Lafora) | Autosomal recessive | LOF |
| **SIK1** | 605705 | Epileptic encephalopathy, early infantile, 30 | Autosomal dominant | n/a |
| **STRADA** | 608626 | Polyhydramnios, megalencephaly, and symptomatic epilepsy | Autosomal recessive | LOF |
| **ARHGEF9** | 300429 | Epileptic encephalopathy, early infantile, 8 | X-linked recessive | LOF |
| **DEPDC5** | 614191 | Epilepsy, familial focal, with variable foci 1 | Autosomal dominant | LOF |
| **GNAO1** | 139311 | Epileptic encephalopathy, early infantile, 17 | Autosomal dominant | n/a |
|  |  | Neurodevelopmental disorder with involuntary movements | Autosomal dominant |  |
| **PLCB1** | 607120 | Epileptic encephalopathy, early infantile, 12 | Autosomal recessive | LOF |
| **TBC1D24** | 613577 | Deafness, autosomal recessive 86 | Autosomal recessive |  |
|  |  | Deafness, autosomal dominant 65 | Autosomal dominant |  |
|  |  | DOOR syndrome | Autosomal recessive | LOF |
|  |  | Epileptic encephalopathy, early infantile, 16 | Autosomal recessive | LOF |
|  |  | Myoclonic epilepsy, infantile, familial | Autosomal recessive |  |
| **CNTNAP2** | 604569 | Cortical dysplasia-focal epilepsy syndrome | n/a | LOF |
|  |  | Pitt-Hopkins like syndrome 1 | n/a |  |
|  |  | {Autism susceptibility 15} | n/a |  |
| **LGI1** | 604619 | Epilepsy, familial temporal lobe, 1 | Autosomal dominant | LOF |
| **PCDH19** | 300460 | Epileptic encephalopathy, early infantile, 9 | X-linked | LOF |
| **RELN** | 600514 | Lissencephaly 2 (Norman-Roberts type) | Autosomal recessive | LOF |
|  |  | {Epilepsy, familial temporal lobe, 7} | Autosomal dominant | LOF |
| **SRPX2** | 300642 | (Rolandic epilepsy, mental retardation, and speech dyspraxia) | n/a | LOF |
| **SZT2** | 615463 | Epileptic encephalopathy, early infantile, 18 | Autosomal recessive | LOF |
| **PAFAH1B1** | 601545 | Lissencephaly | Isolated cases | LOF |
|  |  | Subcortical laminar heterotopia |  |  |
| **CPA6** | 609562 | Epilepsy, familial temporal lobe, 5 | Autosomal dominant;<br>Autosomal recessive | LOF |
|  |  | Febrile seizures, familial, 11 |  |  |
| **CSTB** | 601145 | Epilepsy, progressive myoclonic 1A (Unverricht and Lundborg syndrome) | Autosomal recessive | LOF |
| **NHLRC1** | 608072 | Epilepsy, progressive myoclonic 2B (Lafora) | Autosomal recessive | LOF |
| **SCARB2** | 602257 | Epilepsy, progressive myoclonic 4, with or without renal failure | Autosomal recessive | LOF |
| **ST3GAL5** | 604402 | Salt and pepper developmental regression syndrome | Autosomal recessive | LOF |
| **ST3GAL3** | 138140 | Autosomal Recessive Mental Retardation 12 | Autosomal dominant | LOF |
|  |  | Early Infantile Epileptic Encephalopathy 15 |  |  |
| **KCNC1** | 616187 | Epilepsy, progressive myoclonic 7 | Autosomal dominant | n/a |

**Table S2:** The 40 genes targeted for the Epilepsy Zebrafish Project.

| Human gene | Zebrafish gene | Protein Sequence Reference | % protein identity (GRCz10) | % DIOPT | Homology Score |
| --- | --- | --- | --- | --- | --- |
| *ALDH7A1* | *aldh7a1* | ENSDARP00000108190 | 81 | 75 | 78 |
| *ARHGEF9* | *arhgef9a* | ENSDARP00000118893 | 79 | 100 | 90 |
|  | *arhgef9b* | ENSDARP00000115968 | 84 | 75 | 80 |
| *ARX* | *arxa* | ENSDARP00000075256 | 68 | 75 | 72 |
|  | *arxb* | not identified |  |  |  |
| *CDKL5* | *cdkl5* | ENSDARP00000111280 | 54 | 92 | 73 |
| *CHD2* | *chd2* | ENSDARP00000108411 | 73 | 67 | 70 |
| *CNTNAP2* | *cntnap2a* | (ENSDART00000178326.1) | 71 | 67 | 69 |
|  | *cntnap2b* | ENSDARP00000104097 | 65 | 50 | 58 |
| *CPA6* | *cpa6* | ENSDARP00000096966 | 64 | 83 | 74 |
| *DEPDC5* | *depdc5* | ENSDARP00000098526 | 75 | 58 | 67 |
| *DNM1* | *dnm1a* | ENSDARP00000124266 | 89 | 50 | 70 |
|  | *dnm1b* | ENSDARP00000088100 | 88 | 75 | 82 |
| *EEF1A2* | *eef1a2* | ENSDARP00000010921 | 92 | 92 | 92 |
| *EPM2A* | *epm2a* | ENSDARP00000132560 | 62 | 42 | 52 |
| *GABRA1* | *gabra1* | ENSDARP00000090772 | 84 | 92 | 88 |
| *GABRB3* | *gabrb3* | ENSDARP00000081734 | 73 | 83 | 78 |
| *GABRG2* | *gabrg2* | ENSDARP00000087253 | 83 | 83 | 83 |
| *GNAO1* | *gnao1a* | ENSDARP00000124476 | 90 | 92 | 91 |
|  | *gnao1b* | ENSDARP00000052345 | 84 | 58 | 71 |
| *GOSR2* | *gosr2* | ENSDARP00000069524 | 69 | 83 | 76 |
| *GRIN1* | *grin1a* | ENSDARP00000093144 | 88 | 75 | 82 |
|  | *grin1b* | ENSDARP00000038151 | 88 | 92 | 90 |
| *GRIN2A* | *grin2aa* | ENSDARP00000116766 | 67 | 83 | 75 |
|  | *grin2ab* | not identified |  |  |  |
| *HNRNPU* | *hnrnpua* | ENSDARP00000144487 | 52 | 75 | 64 |
|  | *hnrnpub* | ENSDARP00000112099 | 57 | 83 | 70 |
| *KCNA2* | *kcna2a* | not identified |  |  |  |
|  | *kcna2b* | ENSDARP00000130579 | 92 | 83 | 88 |
| *KCNMA1* | *kcnma1a* | ENSDARP00000118939 | 87 | 67 | 77 |
| *MEF2C* | *mef2ca* | not identified |  |  |  |
|  | *mef2cb* | ENSDARP00000138296 | 74 | 75 | 75 |
| *PAFAH1B1* | *pafah1b1a* | ENSDARP00000042217 | 94 | 92 | 93 |
|  | *pafah1b1b* | ENSDARP00000039257 | 93 | 92 | 92 |
| *PCDH19* | *pcdh19* | ENSDARP00000124001 | 70 | 67 | 68 |
| *PLCB1* | *plcb1* | not identified |  |  |  |
| *PNPO* | *pnpo* | ENSDARP00000011179 | 64 | 75 | 70 |
| *PRICKLE1* | *prickle1a* | ENSDARP00000059513 | 69 | 100 | 85 |
|  | *prickle1b* | not identified |  |  |  |
| *SCN1B* | *scn1ba* | ENSDARP00000079066 | 36 | 100 | 68 |
| *SCN1A* | *scn1laa* | ENSDARP00000138437 | 67 | 67 | 67 |
|  | *scn1lab* | ENSDARP00000125843 | 77 | 58 | 68 |
| *SCN8A* | *scn8aa* | ENSDARP00000024690 | 83 | 83 | 83 |
|  | *scn8ab* | ENSDARP00000126281 | 84 | 75 | 80 |
| *SIK1* | *sik1* | ENSDARP00000077468 | 57 | 75 | 66 |
| *SLC2A1* | *slc2a1a* | ENSDARP00000022579 | 74 | 92 | 83 |
| *SLC6A1* | *slc6a1a* | ENSDARP00000119658 | 73 | 83 | 78 |
|  | *slc6a1b* | ENSDARP00000005281 | 84 | 100 | 92 |
| *SPTAN1* | *spna2* | ENSDARP00000093027 | 90 | 83 | 87 |
| *ST3GAL3* | *st3gal3b* | ENSDARP00000110277 | 63 | 100 | 82 |
| *STRADA* | *strada* | ENSDARP00000115217 | 70 | 67 | 68 |
| *STX1B* | *stx1b* | ENSDARP00000076389 | 97 | 100 | 99 |
| *STXBP1* | *stxbp1a* | ENSDARP00000012776 | 85 | 83 | 84 |
|  | *stxbp1b* | ENSDARP00000026241 | 77 | 67 | 72 |
| *SYNGAP1* | *syngap1a* | ENSDARP00000144044 | 63 | 75 | 69 |
|  | *syngap1b* | ENSDARP00000087797 | 63 | 67 | 65 |
| *TBC1D24* | *tbc1d24* | ENSDARP00000128484 | 55 | 83 | 69 |

**Table S3:**
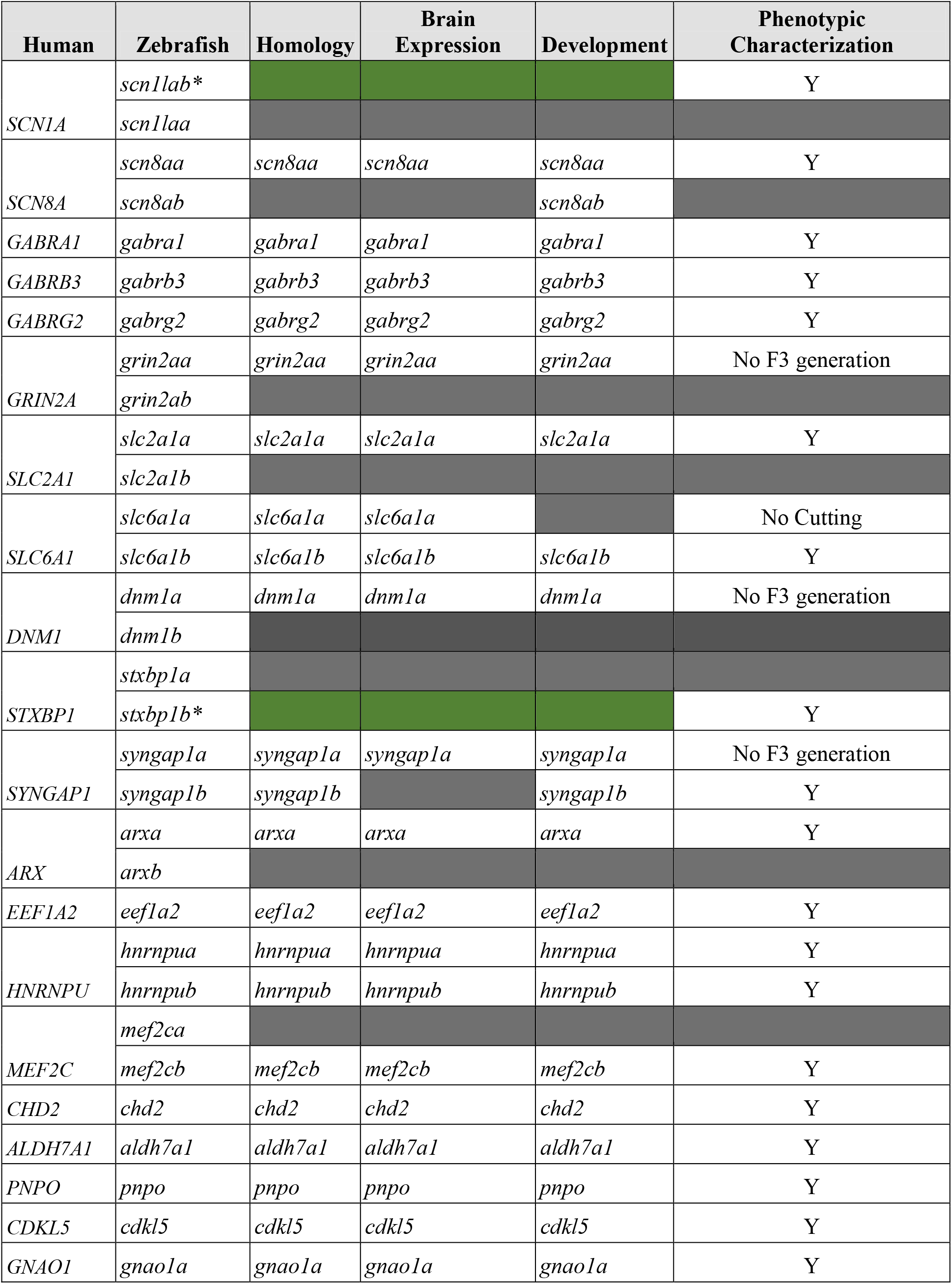
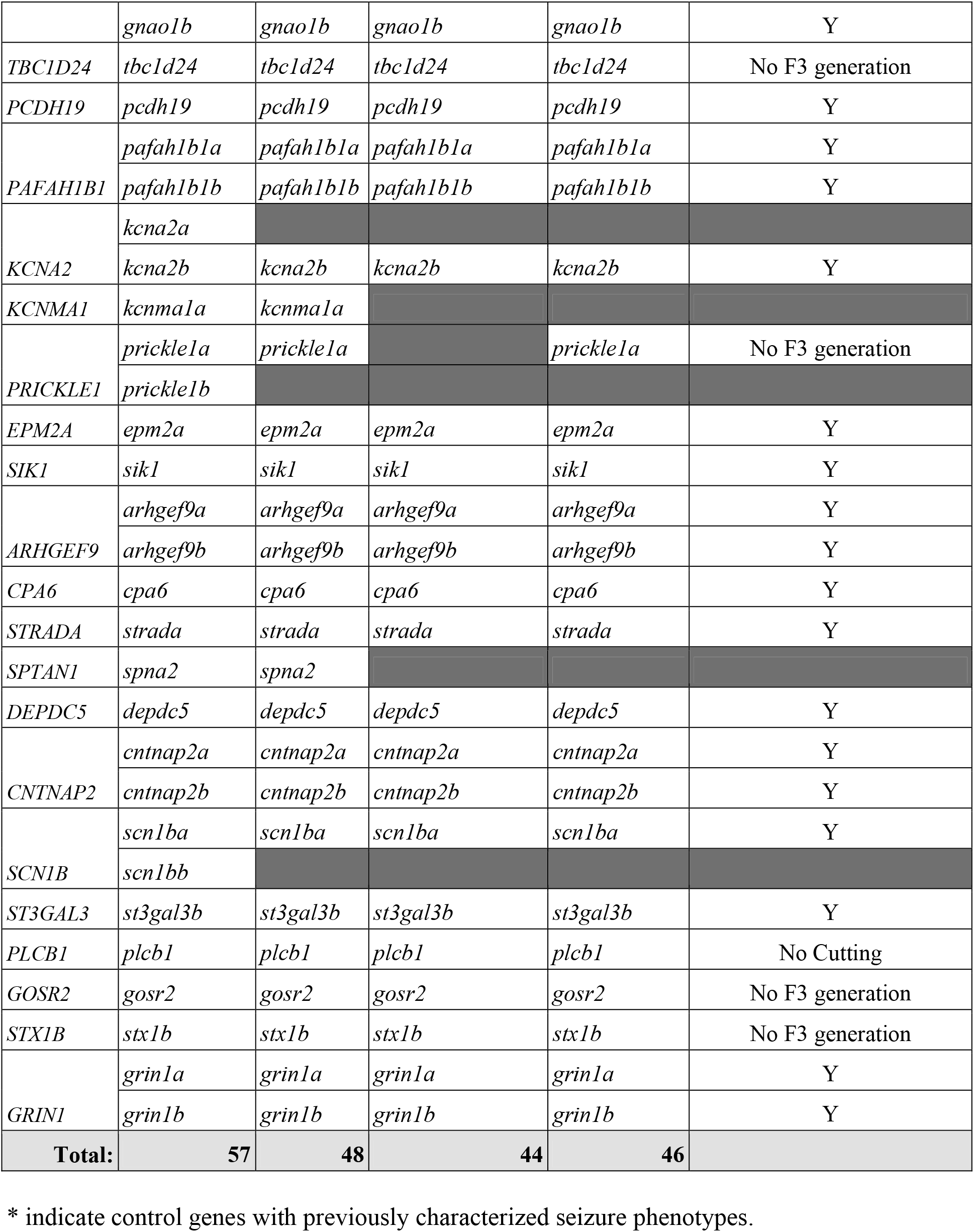
Characterization of zebrafish orthologues for phenotypic characterization.

